# Microbial phenotypic heterogeneity in response to a metabolic toxin: continuous, dynamically shifting distribution of formaldehyde tolerance in *Methylobacterium extorquens* populations

**DOI:** 10.1101/529156

**Authors:** Jessica A. Lee, Siavash Riazi, Shahla Nemati, Jannell V. Bazurto, Andreas E. Vasdekis, Benjamin J. Ridenhour, Christopher H. Remien, Christopher J. Marx

## Abstract

While microbiologists often make the simplifying assumption that genotype determines phenotype in a given environment, it is becoming increasingly apparent that phenotypic heterogeneity (in which one genotype generates multiple phenotypes simultaneously even in a uniform environment) is common in many microbial populations. The importance of phenotypic heterogeneity has been demonstrated in a number of model systems involving binary phenotypic states (e.g., growth/non-growth); however, less is known about systems involving phenotype distributions that are continuous across an environmental gradient, and how those distributions change when the environment changes. Here, we describe a novel instance of phenotypic diversity in tolerance to a metabolic toxin within wild-type populations of *Methylobacterium extorquens,* a ubiquitous phyllosphere methylotroph capable of growing on the methanol periodically released from plant leaves. The first intermediate in methanol metabolism is formaldehyde, a potent cellular toxin that is lethal in high concentrations. We have found that at moderate concentrations, formaldehyde tolerance in *M. extorquens* is heterogeneous, with a cell’s minimum tolerance level ranging between 0 mM and 8 mM. Tolerant cells have a distinct gene expression profile from non-tolerant cells. This form of heterogeneity is continuous in terms of threshold (the formaldehyde concentration where growth ceases), yet binary in outcome (at a given formaldehyde concentration, cells either grow normally or die, with no intermediate phenotype), and it is not associated with any detectable genetic mutations. Moreover, tolerance distributions within the population are dynamic, changing over time in response to growth conditions. We characterized this phenomenon using bulk liquid culture experiments, colony growth tracking, flow cytometry, single-cell time-lapse microscopy, transcriptomics, and genome resequencing. Finally, we used mathematical modeling to better understand the processes by which cells change phenotype, and found evidence for both stochastic, bidirectional phenotypic diversification and responsive, directed phenotypic shifts, depending on the growth substrate and the presence of toxin.

**Author Summary:** Scientists tend to appreciate microbes for their simplicity and predictability: a population of genetically identical cells inhabiting a uniform environment is expected to behave in a uniform way. However, counter-examples to this assumption are frequently being discovered, forcing a re-examination of the relationship between genotype and phenotype. In most such examples, bacterial cells are found to split into two discrete populations, for instance growing and non-growing. Here, we report the discovery of a novel example of microbial phenotypic heterogeneity in which cells are distributed along a gradient of phenotypes, ranging from low to high tolerance of a toxic chemical. Furthermore, we demonstrate that the distribution of phenotypes changes in different growth conditions, and we use mathematical modeling to show that cells may change their phenotype either randomly or in a particular direction in response to the environment. Our work expands our understanding of how a bacterial cell’s genome, family history, and environment all contribute to its behavior, with implications for the diverse situations in which we care to understand the growth of any single-celled populations.

## Introduction

Microbes are individuals. Even in seemingly simple unicellular organisms, phenotype is not always the straightforward product of genotype and environment; cells with identical genotypes in identical environments may nonetheless demonstrate cell-to-cell diversity in the expression of any of a number of traits. Frequently overlooked in everyday microbiology experiments, the phenomenon of cell-to-cell phenotypic heterogeneity has drawn increasing attention in recent decades both from a systems biology perspective and from an evolutionary perspective, as well as for its consequences to applied fields such as medicine (e.g., antibiotic persistence [1]; cancer cell drug tolerance [2, 3]) and biological engineering [4].

Some forms of population heterogeneity might be considered trivial: molecular interactions within cells are inherently noisy. All genes might be expected to be expressed at slightly different levels among different cells [5–7], and historical contingency (e.g., pole age, asymmetrical division of macromolecules) can also create inherent diversity within microbial populations, independent of signals from the environment [8–10]. Naturally, evolution imposes some pressure on organisms to limit the noise in pathways that are essential for life [11]; what is more remarkable is that some pathways seem to be selected for increased noise, and in many cases that noise is further amplified by feedback circuits, enabling a population to split into different phenotypes. Specifically, genes involved in stress response and in metabolism have been found to show higher heterogeneity in expression than those in other pathways [12], and many of the well-understood examples of binary phenotypes involve stress response (antibiotic persistence [13]; sporulation [14]), or carbon transport and use (lactose utilization in *E. coli* [15]; diauxic switch in *S. cerevisiae* [16, 17]). While some forms of phenotypic noise may have little fitness impact, phenotypic heterogeneity involving binary phenotypes is argued in many cases to offer an evolutionary advantage, as a strategy to facilitate diversifying bet-hedging or division of labor [12, 18]. In many cases, the genetic basis of phenotypic differentiation is known, and laboratory evolution studies have demonstrated how populations can evolve the timing or frequency of that differentiation in response to environmental selection [19, 20]. Moreover, it is argued that phenotypic heterogeneity influences the rate of genotypic evolution either by creating an “epigenetic load” that contributes to extinction [21], or by allowing populations to adapt faster to changing environments and by increasing the opportunities for mutations to arise and reach fixation [22, 23].

Besides cases of binary phenotypes or modest phenotypic noise around a population mean, a third possibility involves phenotypes that fall along a continuous gradient. Fewer such phenomena have been described [12], but some examples include populations in which cells have a range of levels of stress tolerance: for instance, a genetically chloramphenicol (Cm)-resistant *E. coli* strain exhibits a wide, continuously-varying distribution of maximum concentrations of Cm that individual cells can tolerate. The population splits into growing and non-growing subpopulations in the presence of Cm, and the proportion of growing cells varies according to the environmental Cm concentration [24]. This effect is mediated by a positive-feedback interaction between intracellular chloramphenicol acetyltransferase (CAT) activity and innate cell growth; thus, the ultimate fate of an individual cell depends on its initial internal CAT activity, which can vary continuously among cells. If such a population were to experience shifts in environmental Cm concentrations, the population distribution of per-cell CAT activity levels could presumably shift as a consequence of either cellular responses (e.g., CAT upregulation), or simply by selection against sensitive cells; however, this has not been described. Analogous population-level phenotype distribution shifts have been explored only through mathematical modeling, for instance in the case of human cancer cells exhibiting a gradient of tolerances to cytotoxic drugs [25]. In this case, it is assumed that random epimutations result in small phenotypic variations in drug tolerance, and that drug exposure leads to selection upon that diversity. However, experimental work would be necessary to verify whether modeling accurately predicts cell population dynamics, or whether other processes—such as those in which cells sense drug concentrations and respond with phenotypic shifts—might also play a role. Examples of phenotypic heterogeneity in which a population with a continuous phenotype distribution interacts dynamically with the environment to undergo dramatic shifts in population distributions, pose complex—yet unanswered—questions about the importance of phenotypic heterogeneity to population-level fitness in diverse environments.

Here we present a novel example of a continuously-distributed threshold phenotype in an environmentally relevant microorganism, *Methylobacterium extorquens*, and describe the dynamics of that phenotype distribution in response to shifts in its growth environment. *M. extorquens*, a species of *Alphaproteobacteria* found ubiquitously on plant leaves, is a methylotroph: it can grow on reduced single-carbon compounds such as methanol, which is emitted from leaves through the activity of plant pectin methylesterases [26]. The first intermediate in methanol metabolism is the potent toxin formaldehyde [27, 28], an electrophile that can cause cellular damage through its reactions with macromolecules, and can be lethal to microorganisms [29]. In *M. extorquens*, formaldehyde is produced in the periplasm through the oxidation of methanol by methanol dehydrogenase (MDH); in the cytoplasm it is then oxidized via a tetrahydromethanopterin (H_4_MPT)-dependent pathway. *M. extorquens* therefore exists in a tension between the two goals of rapid substrate utilization and prevention of the accumulation of a toxic intermediate. The importance of formaldehyde oxidation for single-carbon metabolism has been demonstrated by the inability of H_4_MPT-pathway mutants to grow in the presence of methanol when they possess a functional MDH [30]. However, although MDH activity is constitutive, it has also been observed that downstream single-carbon assimilation pathways are up-regulated only in the presence of methanol; the consequence is that when cells previously grown on a multi-carbon substrate are first exposed to methanol, formaldehyde production initially outpaces consumption so drastically that it is excreted into the medium [27]. This is just one example of many potential situations in which cellular formaldehyde accumulation poses a threat to *M. extorquens*, and yet we know very little about how the species copes with formaldehyde toxicity.

As an initial step toward understanding the effect of formaldehyde toxicity on *M. extorquens*, we conducted time-kill experiments in which we exposed cells to a range of formaldehyde concentrations in batch liquid culture and monitored their viability over time. To our knowledge, previous research on formaldehyde toxicity in *M. extorquens* has consisted only of single-timepoint shock experiments, or a pulse of methanol added to succinate-grown cultures to cause cellular formaldehyde production [30]. We hoped that making time-resolved measurements in extreme formaldehyde conditions would shed light on both large-scale patterns of toxicity and on cell behavior near the minimum bactericidal concentration (MBC). Indeed, quite unexpectedly, we found cell-to-cell variation in the MBC within isogenic populations of *M. extorquens*. We confirmed this to be a phenomenon of phenotypic diversity at the single-cell level, and investigated its dynamic population-level consequences, using a combination of liquid culture experiments, colony growth tracking, flow cytometry, single-cell time-lapse imaging, transcriptomics, genome resequencing, and mathematical modeling.

## Results

### Formaldehyde-induced death occurs at an exponential, concentration-dependent rate above 5 mM

In order to better understand the physiological effects of the toxic metabolite formaldehyde on *M. extorquens*, we initially conducted a series of experiments in which we added formaldehyde to methanol growth medium and assessed the relationship between toxin concentration and mortality in well-mixed liquid batch culture. We found that concentrations ≥5 mM elicited loss of viability (as measured in colony-forming units, CFUs) at an exponential rate, and the rate of death increased with increasing formaldehyde concentration (Fig. 1). (The full data shown in all figures are available in Supporting Information, Data S1.) A concentration of 15 mM was sufficient to eliminate all detectable viable cells within 1.5 hours (approximately half of one generation time). These time-kill curves indicate that formaldehyde-induced death can be modeled as a single-hit process [31], as is often observed for other multi-target lethal agents such as radiation and heat [32], and some bactericidal antibiotics [33, 34]. Given that the precise mechanism of formaldehyde-induced mortality remains unknown but likely involves multiple cellular targets, it is noteworthy that formaldehyde-induced death over time did not seem to involve substantial “shoulders” or “tails” as is sometimes observed with other agents [34], and that we observed no saturation of death rate at the concentrations tested. In analogy to bactericidal antibiotics, our results suggested that the minimum bactericidal concentration (MBC) of formaldehyde for *M. extorquens* is 5 mM.

**Fig 1.**
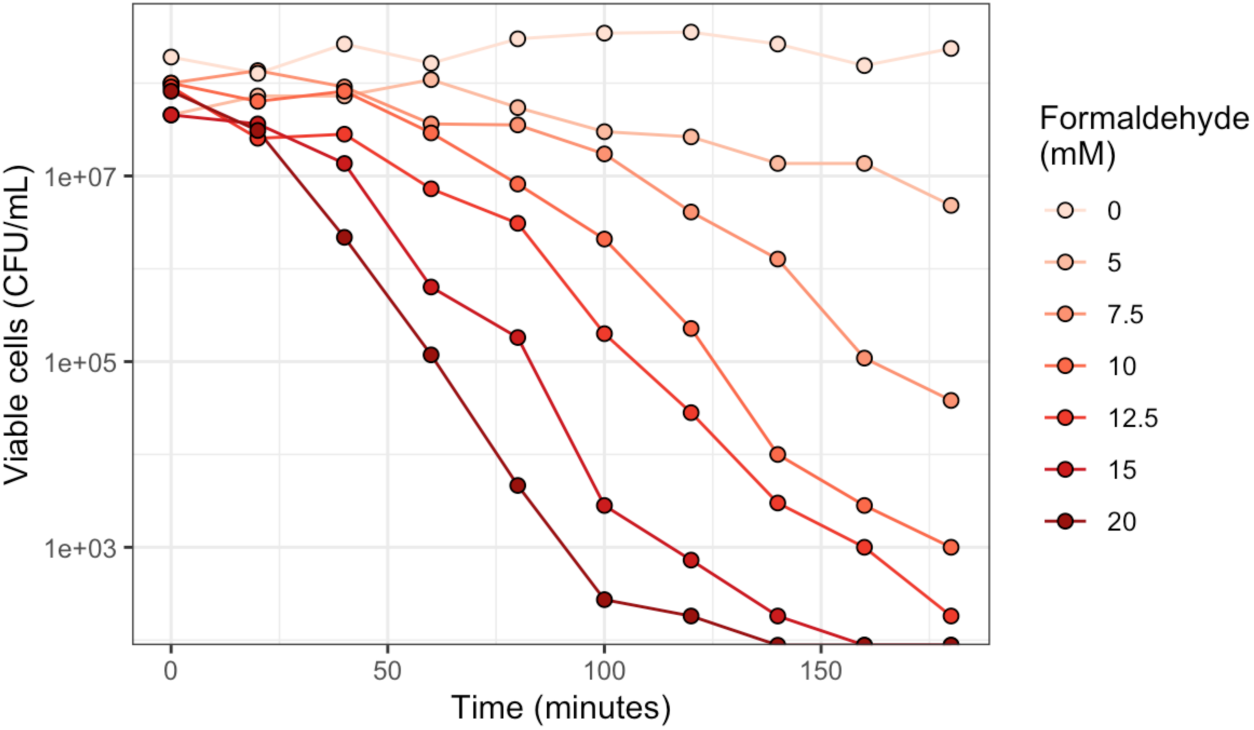
Formaldehyde kills *M. extorquens* at an exponential, concentration-dependent rate. Formaldehyde was added at the indicated concentrations to liquid cultures of *M. extorquens* cells growing in minimal medium with methanol, and abundance of viable cells was measured as colony-forming units (CFU) over time. Note that negligible growth is expected to have occurred during the course of this experiment, as the 180-minute duration was less than one generation (∼3.5 hrs) for *M. extorquens* in these conditions. The original data shown in this and all other figures are available in Supporting Information file Data S1.

### Exposure to moderate formaldehyde concentrations shifts population trajectory

Follow-up experiments on longer timescales (3–4 days) revealed that assessing MBC was in fact not straightforward: lower concentrations had an effect on *M. extorquens* growth as well. Exposure to formaldehyde concentrations between 3 and 5 mM allowed normal growth of *M. extorquens* as measured by optical density (OD_600_), but only after an apparent lag time of several hours to days. Higher formaldehyde concentrations induced longer lags, but growth subsequently resumed, and formaldehyde concentration had no effect on growth rate (for 0, 3, 4, and 5 mM respectively, specific growth rates (*r*) were 0.212±0.038, 0.208±0.045, 0.238±0.002, and 0.230±0.018 h^-1^, where ± indicates 95% confidence interval; *p*=0.237 for the effect of treatment group on growth rate by ANOVA) (Fig. S1). To better understand the apparent lag in these conditions, we measured cell viability over time during formaldehyde exposure experiments. CFU measurements revealed that the apparent “lag time” was in fact not a lag, but rather a period of exponentially decreasing cell counts followed by an abrupt transition to increasing counts (Fig. 2, round gray symbols and line). These dynamics repeated themselves across multiple biological replicates at multiple formaldehyde concentrations, with remarkable consistency in the timing of the transition between population decrease and increase. At a concentration of 4 mM formaldehyde, this transition occurred at approximately 20 hours. Upon recovery, the increase in CFUs was exponential and the rate was nearly equivalent from growth rates on methanol in the absence of external formaldehyde (*r*=0.215±0.001 h^-1^ for 0 mM; *r*=0.202±0.030 h^-1^ for 4 mM; *p*=0.063 by Welch’s t-test) (Fig. 2).

**Fig 2.**
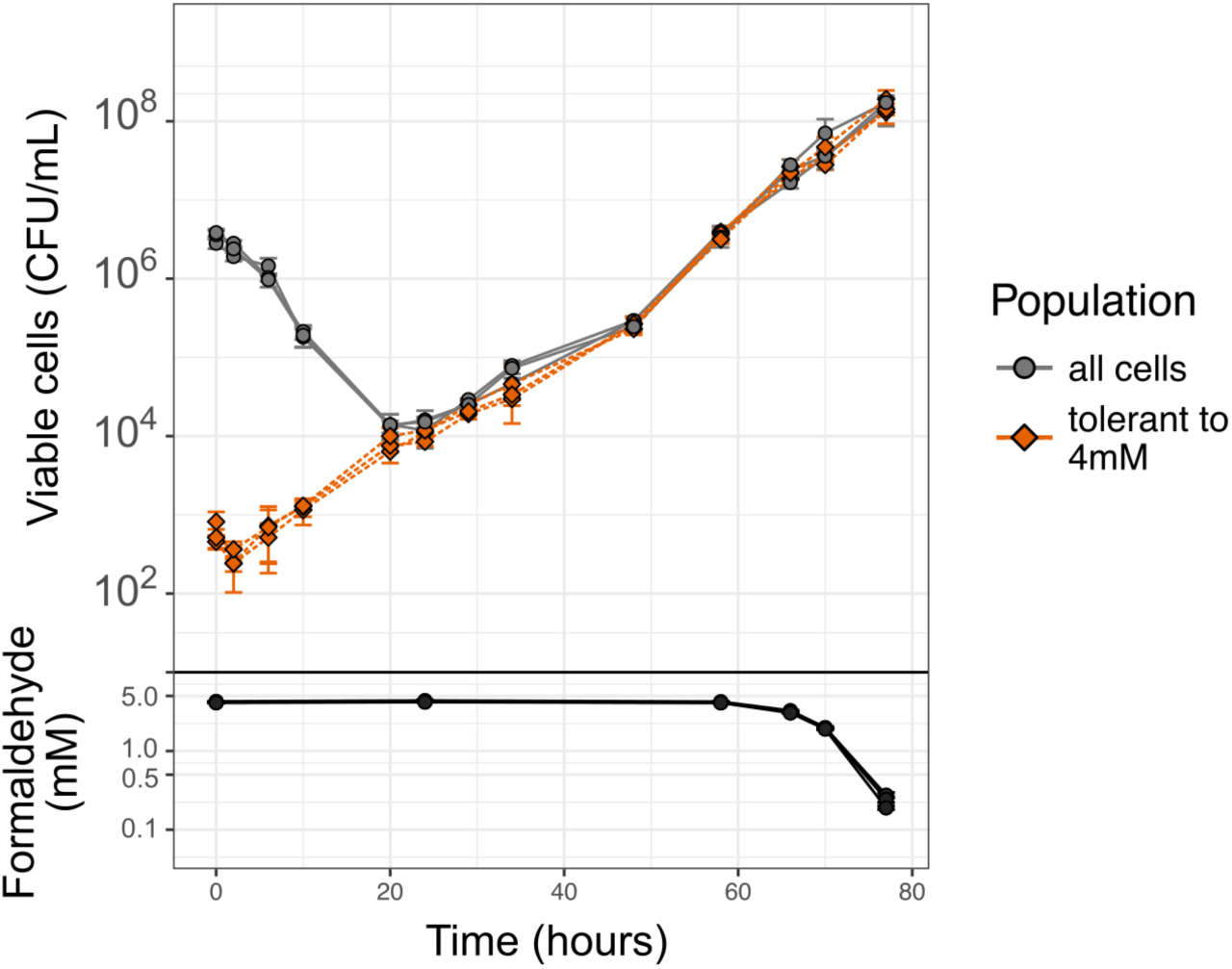
Re-growth of *M. extorquens* after population decline in the presence of formaldehyde is due to a pre-existing sub-population of formaldehyde-tolerant cells. Stationary-phase cells were inoculated into fresh medium containing methanol and 4 mM formaldehyde. The abundance of viable cells in the two different populations was assessed over time by removing and washing cells, then plating onto both permissive medium (without formaldehyde: “all cells”) and selective medium (with 4 mM formaldehyde: “tolerant to 4 mM”). CFU = colony-forming units. Each line represents one biological replicate; error bars show the standard deviation of three replicate platings. Formaldehyde in the liquid medium during the incubation period was measured by a colorimetric assay on subsamples after removing cells by centrifugation.

We explored several hypotheses to explain the abrupt decrease-then-increase transition we observed during formaldehyde exposure. These included: 1) consumption of formaldehyde by the cells; 2) the existence of formaldehyde-resistant genetic mutants; 3) phenotypic changes in the plating efficiency of cells due to formaldehyde-induced damage and its repair; 4) the existence of a subpopulation of phenotypically (but not genetically) formaldehyde-tolerant cells.

### Populations recover despite high formaldehyde concentrations

One explanation for the abrupt change from population decline to population increase might be a change in environmental conditions. For instance, cooperative behavior to metabolize a toxin to sublethal levels can result in population rescue even if not all cells are tolerant, as has been observed with antibiotics [35]. We therefore investigated the possibility that formaldehyde consumption had reduced the toxin concentration in the medium. In formaldehyde exposure experiments with 4 mM formaldehyde, we monitored formaldehyde concentrations in the medium for 80 hours, a period that encompassed the death and regrowth phases. Although *M. extorquens* is capable of metabolizing formaldehyde, we found that a measurable decrease in the concentration of the toxin only occurred at the very end of the growth period in batch culture, more than 40 hours after the decrease-then-increase transition (Fig. 2, black symbols and line). A change in the toxicity of the environment was therefore not responsible for the change in population trajectory.

### Genetic mutations are not responsible for population recovery

The next most parsimonious explanation was that the observed growth was due to a small pre-existing subpopulation of formaldehyde-tolerant mutants whose existence became apparent only after the death of the sensitive majority. To assess this possibility, we grew cells in the presence of 4 mM formaldehyde for 80 hours as described above, then subcultured them into fresh medium without formaldehyde for 6 generations (using succinate as the growth substrate), followed by another subculture into formaldehyde-containing medium. The population decrease-then-increase dynamics were recapitulated, indicating that formaldehyde tolerance was not transmitted in a manner consistent with genetic heritability, and that the descendants of cells tolerant to 4 mM formaldehyde were re-sensitized in its absence (see “Transitions between tolerance phenotypes,” below, for more detail).

In addition, we prepared genomic DNA from cells that had grown in the presence of formaldehyde (the 80-hour timepoint of a 4 mM formaldehyde exposure experiment), and used this for whole-genome resequencing. The resequenced genome was compared to the published genome sequence of wild-type *M. extorquens* PA1 and to that of an *M. extorquens* population grown without formaldehyde; no evidence was found for SNPs, deletions, insertions, or gene duplications in the formaldehyde-selected population. These sequencing data, along with the instability of the formaldehyde tolerance phenotype, indicate that the heterogeneous formaldehyde tolerance we have observed in *M. extorquens* is not due to genetic mutations.

### Population recovery is not due to changes in plating efficiency

Having ruled out both environmental change and genetic mutations as explanations for the sharp transition between population decrease and increase during formaldehyde exposure, we pursued the possibility that the observed dynamics might be due to a change in phenotype. As our evidence for population dynamics thus far was based on counts of colony-forming units on agar medium, one possibility was that the phenotypic change might be related to the ability of cells to form colonies. In this scenario, formaldehyde exposure might cause cellular damage resulting in a decrease in cells’ ability to form colonies and their entry into a viable-but-not-culturable (VBNC) state [36], and the inflection point at 20 hours represented the beginning of recovery of these same cells, rather than turnover of the population.

To investigate whether there had been population turnover, we carried out a cell proliferation assay, which enabled us to interrogate single-cell growth dynamics in liquid culture without plating (as in [37]). We used a nontoxic fluorescent membrane linker dye to stain the population prior to conducting a 4 mM formaldehyde exposure experiment as described above, and then assessed the trajectory of per-cell fluorescence over time by flow cytometry. In exponentially growing populations, the membrane dye of each parent cell was divided between its two daughter cells, and per-cell fluorescence decreased uniformly across the population as the number of cells increased (Fig. 3, upper left). In non-growing populations, as when *M. extorquens* was exposed to 20 mM formaldehyde, membrane dye remained in the cells and did not fade, and per-cell fluorescence remained the same (Fig. 3, lower left). However, in an *M. extorquens* population treated with 4 mM formaldehyde, two populations were evident: one group of cells that retained the same membrane fluorescence throughout the experiment, and another group of cells that increased in number and decreased in per-cell membrane fluorescence, indicating normal growth in the presence of formaldehyde (Fig. 3, right). At 4 mM, the growing population was observable only after ∼37 hours of incubation, consistent with a population that began in extremely low abundance relative to the non-growing population. These data indicate that there was population turnover, and that the increase in viable cell counts was in fact due to the growth of a tolerant, rare subpopulation of cells that remained active in the presence of formaldehyde.

**Fig 3.**
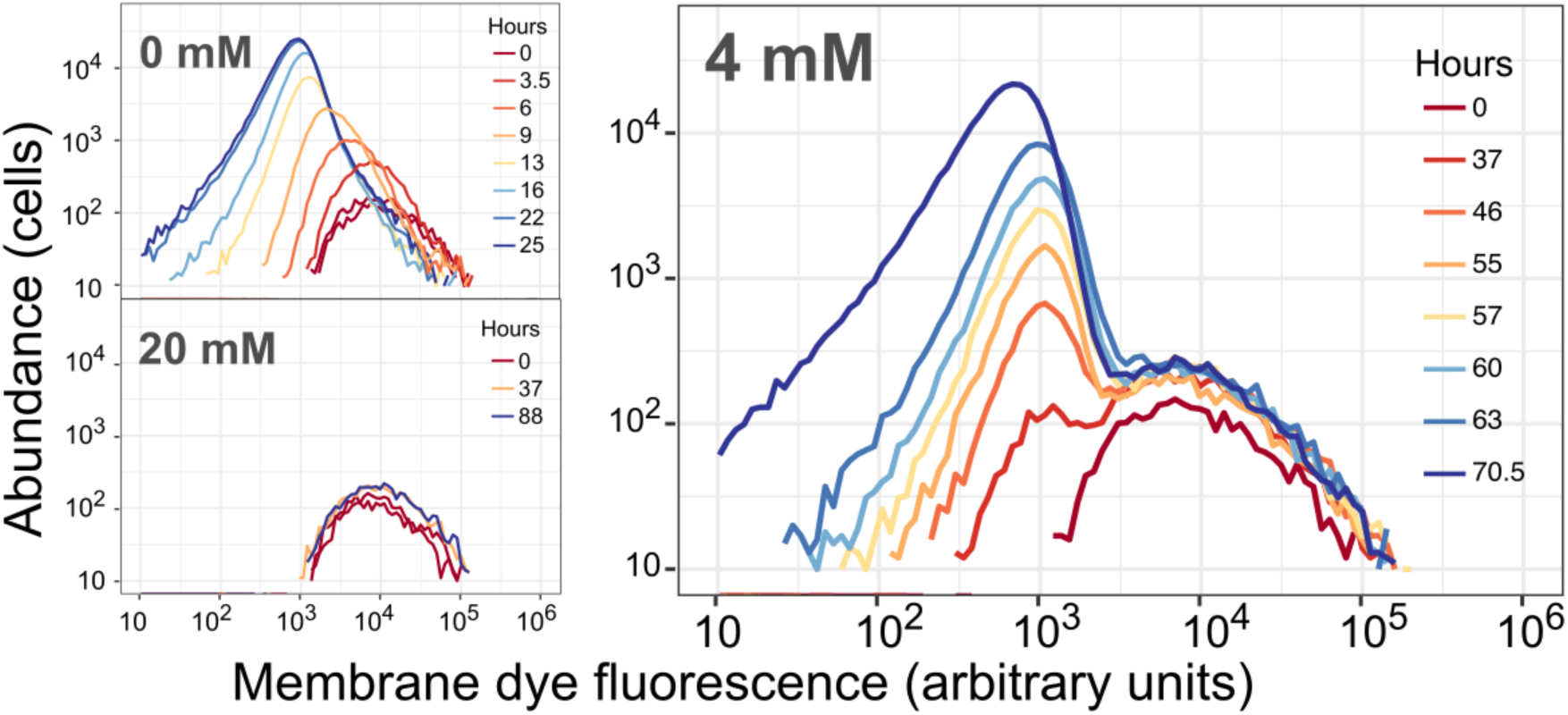
Cell proliferation assay shows dynamics consistent with the coexistence of both growing and non-growing subpopulations, with no turnover between the two. Cells were stained with PKH67 fluorescent membrane dye, then allowed to grow in minimal medium with methanol and either 0, 4, or 20 mM formaldehyde. Histograms show per-cell fluorescence of the cells (events measured by flow cytometry) present in 30 µL of culture at each timepoint; colors denote the time of sampling in hours (note that different color scales are used in different panels). Top left: without formaldehyde, all cells underwent doubling, diluting their membrane fluorescence so that the median fluorescence decreased as population increased. Bottom left: at high concentrations of formaldehyde, no cells grew, leaving per-cell fluorescence unchanged. Right: in the presence of 4 mM formaldehyde, most cells did not grow, but a few did; consequently, a small growing population with lower per-cell fluorescence became detectable at 37 hours and continued to increase in abundance thereafter. Results of experiments conducted at other formaldehyde concentrations are shown in Fig. S5. Flow cytometry data are provided in Supporting Information, Data S3.

### Phenotypically tolerant subpopulation is present prior to formaldehyde exposure

Given our evidence against genetic mutants, a fourth hypothesis was that the observed growth was due to a small subpopulation of cells that were phenotypically (but not genetically) highly tolerant to formaldehyde. Stress-tolerant individuals may arise in microbial populations stochastically, or they may do so in response to an environmental signal [38]. To investigate whether the hypothesized formaldehyde-tolerant cells were present in the original population or were induced during the course of formaldehyde exposure, we monitored the abundance of formaldehyde-tolerant CFU over time. Specifically, we repeated the 4 mM formaldehyde exposure experiment and plated the cells harvested at each timepoint on selective agar culture medium (4 mM formaldehyde, allowing the growth of only tolerant cells) and permissive medium (without formaldehyde, to enumerate all cells). We found that at the beginning of the experiment, the *M. extorquens* population already contained a detectable subpopulation of cells that were able to form colonies in the presence of 4 mM formaldehyde. This subpopulation comprised only a small portion of the total number of cells (a frequency of ∼10^-4^ in the total population of ∼2×10^6^ cells) (Fig. 2, orange diamond symbols and lines). And while the total abundance of cells decreased at an exponential rate between 0 and 20 hours, the formaldehyde-tolerant subpopulation increased at a constant rate for nearly the entire course of the experiment, such that after 20 hours the population was dominated by cells tolerant to 4 mM formaldehyde.

### Formaldehyde-tolerant subpopulation shows no evidence of formaldehyde-induced cell damage

The fact that the tolerant subpopulation increased at a rate commensurate with standard growth in formaldehyde-free media suggests that these tolerant cells may have escaped formaldehyde-induced mortality and any damage that might impede or delay growth. However, some stress-tolerant phenotypes, such as persister cells, are characterized by slow growth or even absence of growth [13]. We therefore looked more closely at the growth phenotypes of the tolerant subpopulation: specifically, whether they differed in their rate of colony growth, or in the time required to initiate a colony, relative to normal unstressed cells. We repeated the 4 mM formaldehyde exposure experiment described above, this time using time-lapse imaging to monitor the growth of colonies resulting from timepoint samples plated on both selective and permissive plates. Plates were incubated on a flatbed photo scanner and images captured hourly and processed to extract per-colony statistics (Fig. S2), in a manner similar to previous studies investigating cell damage and dormancy at the single-cell level [39–41].

We found that within the total population, which consisted of primarily of sensitive cells at the beginning of the experiment (tolerant cells were present at a frequency of only 10^-4^), formaldehyde exposure prior to plating did indeed induce a lag in colony formation, such that each hour of formaldehyde exposure led to an increase in colony appearance time (the time necessary to form a detectable colony) of approximately 4.80 hours (r^2^=0.766 by linear regression, *p*<0.005) (Fig. 4). This pattern held true for the first 16 hours of exposure, until most of the sensitive cells lost viability entirely. At 16 hours, we observed a clear bimodality on the permissive plates, with approximately half of the cells appearing late and the other half appearing early, and the abundance of the early colonies on the permissive plates was consistent with the abundance of the tolerant subpopulation (all colonies on the selective plates). However, on the selective plates, we observed no relationship between appearance time and the length of formaldehyde exposure prior to plating (slope=-0.091 hours lag per hour exposure, r^2^=0.0639, *p*<0.005), supporting the hypothesis that the tolerant subpopulation is in a different physiological state that does not experience formaldehyde damage in the same way as the sensitive subpopulation. Furthermore, we observed no trend in average growth rate due to formaldehyde exposure time (slope=-2.83×10^-4^ hours lag per hour exposure, r^2^=6.62×10^-4^, *p*=0.123), although colonies from samples that had been exposed for longer had more varied growth rates (there was a positive linear relationship between exposure time and the log of the median average deviation of growth rate, *p*<0.05, for both permissive and selective plates).

**Figure 4.**
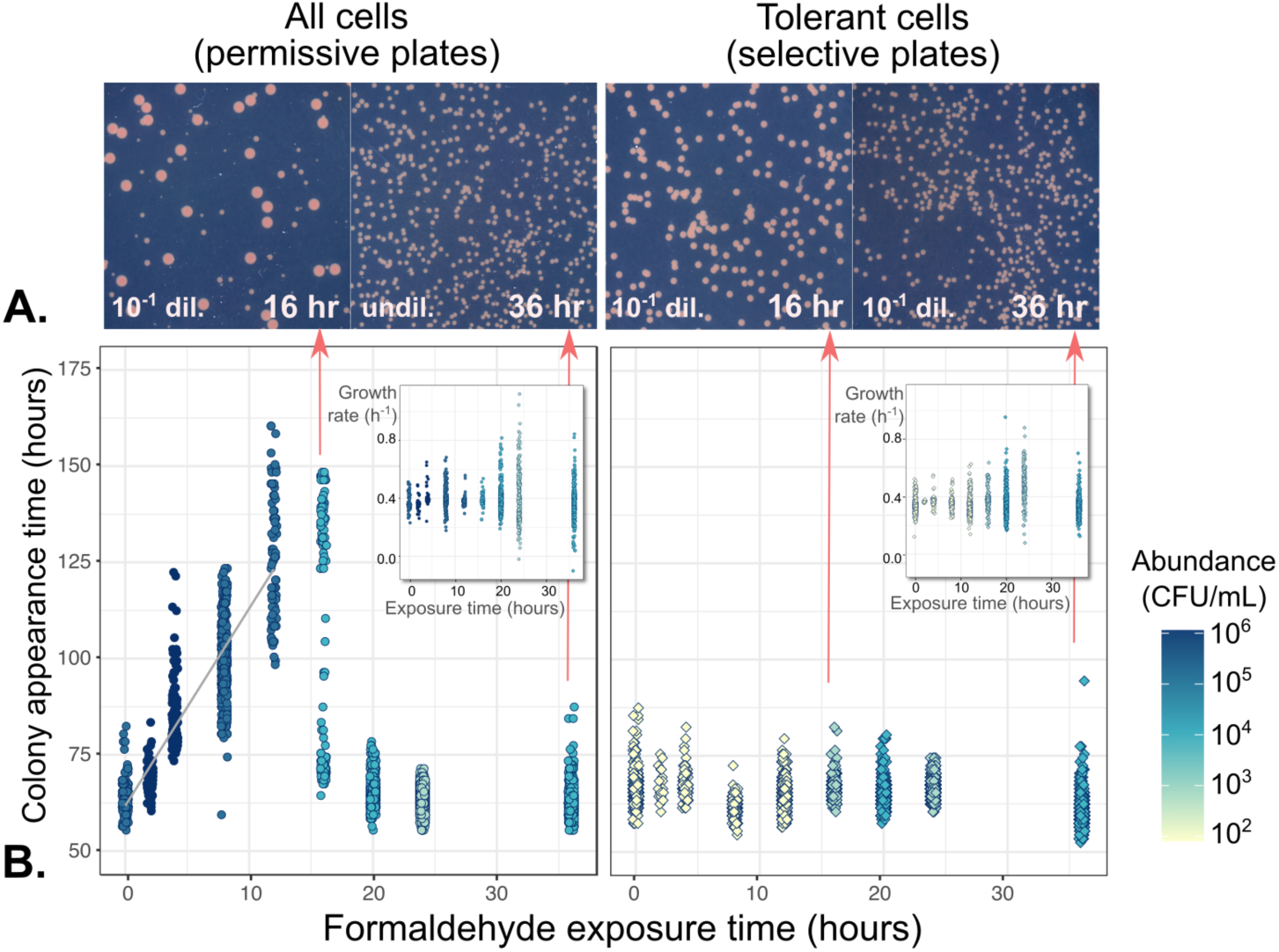
Cell damage by formaldehyde results in delayed colony appearance for the majority of cells, but not for the tolerant subpopulation. Cells from a formaldehyde exposure experiment (liquid MPIPES medium with 4 mM formaldehyde) were sampled at 2- to 4-hour intervals, washed, and plated onto both permissive medium (no formaldehyde, allowing the growth of all cells) and selective medium (4 mM formaldehyde, allowing the growth of only the tolerant subpopulation). A) Images of colonies on plates. Colony size heterogeneity was evident only on permissive medium with cultures exposed to formaldehyde for 16 hours, consistent with a population containing both sensitive cells that formed colonies late due to formaldehyde-induced damage (small colonies) and tolerant cells that formed colonies early (large colonies). All images are shown at the same magnification level; *dil*=dilution factor prior to plating. B) Relationship between formaldehyde exposure and colony growth characteristics. Shading indicates abundance of colony-forming units in each population (see Fig. 2); samples were diluted prior to plating for an average of 500 colonies per plate. Left panel: gray line shows linear regression of appearance time on exposure time for the first 12 hours. Every hour of exposure to formaldehyde led to a ∼4.8-hour delay in colony appearance time among sensitive cells. At 16 hours, the population consisted of both damaged and tolerant cells; after 20 hours, all cells were tolerant due to the death of the damaged cells. Right panel: among tolerant cells, formaldehyde exposure had no effect on appearance time. Insets: exposure time affected only the variability among colony growth rates, but not their median.

### Gene expression is significantly different between tolerant and sensitive populations

To determine whether the different physiology of tolerant versus sensitive subpopulations might be attributed to differences in gene expression, we conducted transcriptomic sequencing on three sets of samples from a 4 mM formaldehyde exposure experiment. To distinguish features associated with tolerance from those due to formaldehyde stress, we examined i) an exponentially-growing population of cells not exposed to formaldehyde stress, ii) an exponentially-growing tolerant subpopulation (cultures exposed to formaldehyde for 64 hours), and iii) sensitive cells that were declining in viability due to formaldehyde toxicity (cultures exposed to formaldehyde for 4 hours). For both formaldehyde-exposed populations, we calculated differential gene expression relative to growing cells not exposed to formaldehyde stress (the majority of which are sensitive).

Sensitive cells experiencing formaldehyde stress showed extensive, global changes in gene transcription: out of a genome with 4,829 loci, 2,591 (53.6%) were significantly up- or down-regulated (greater than 1.0 log_2_ fold change with an adjusted *p*-value of <0.05) relative to the unstressed population. In contrast, the tolerant subpopulation growing in the presence of formaldehyde had only 23 differentially expressed genes, most of which were up-regulated (Fig. 5, Table 1). None of these genes showed a significant change in the same direction in the sensitive stressed population, and the majority (17) had a significant change in the opposite direction. This suggests that tolerance and stress response are distinct, and often opposing, phenotypes. The 23 genes represented 6 distinct clusters in the genome, each cluster consisting of one or a few adjacent operons.

**Fig 5.**
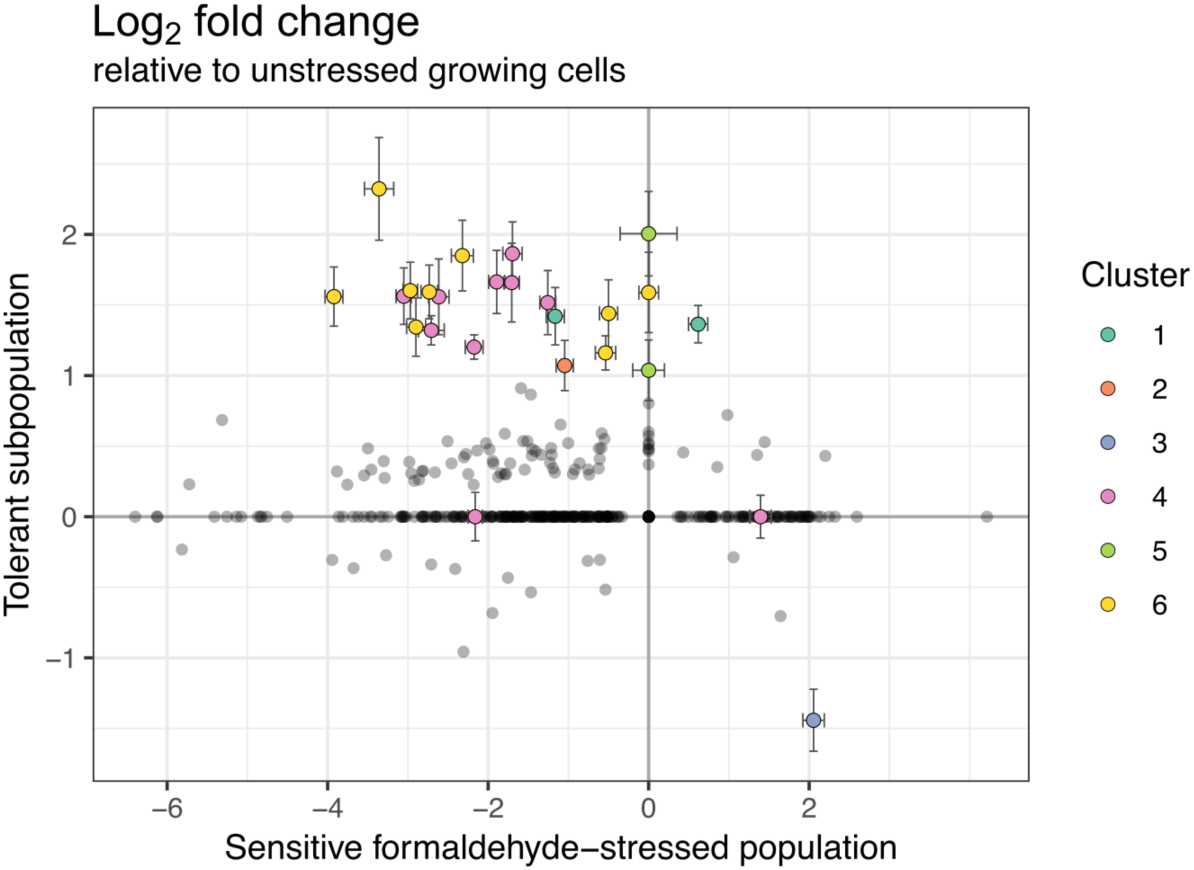
Gene expression in the formaldehyde-tolerant subpopulation is distinct from that of both unstressed and formaldehyde-stressed sensitive populations. RNA sequencing was carried out on cells from a 4 mM formaldehyde exposure experiment harvested at 4 hours (sensitive cells losing viability due to formaldehyde toxicity), and 64 hours (selected formaldehyde-tolerant cells in exponential growth in the presence of formaldehyde). Log_2_ fold change was calculated relative to an unstressed population growing in the absence of formaldehyde. Each circle represents the expression of one gene in the tolerant population (y-axis) versus in the sensitive population (x-axis). Colored circles are genes with >1.0 log_2_ fold change and *p*_adj_>0.05 in the tolerant population, as well as genes belonging to the same clusters in the genome (23 genes total, 6 clusters); these are described in Table 1. Error bars indicate standard error for the log fold change estimate. Most genes that were differentially expressed in the tolerant population showed opposite expression patterns in the stressed population.

**Table 1.**
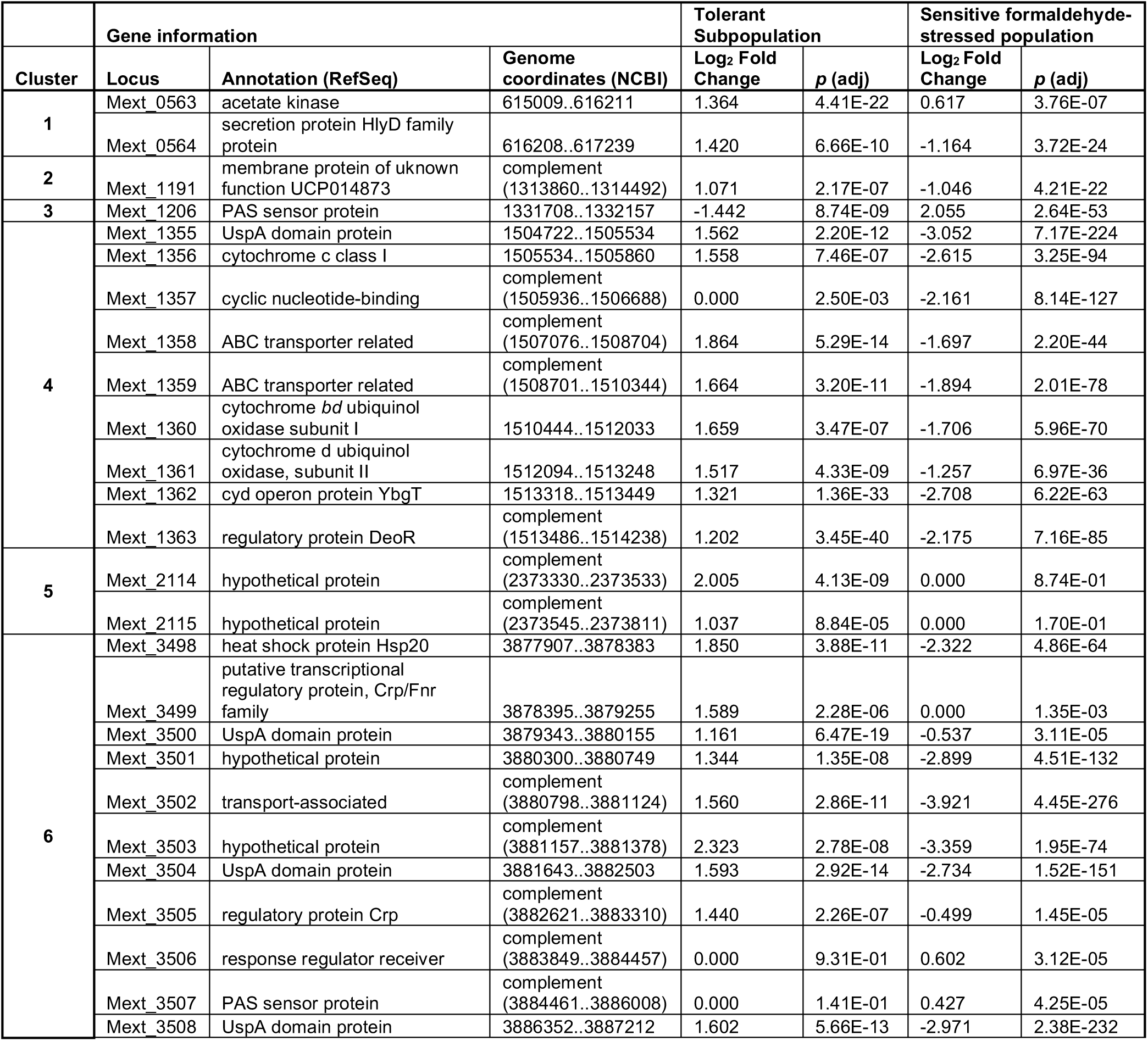
Gene regions differentially expressed in the formaldehyde-tolerant subpopulation. Cells were harvested in exponential growth after selection on methanol with 4 mM formaldehyde for 64 hours. Shown here are all the genes with log_2_ fold change > 1.0 and adjusted *p*-value > 0.05 relative to a population growing on methanol with no formaldehyde. There were three genes located within the same clusters in the genome that had no change in expression; these are also included. Expression levels in the tolerant subpopulation are shown alongside expression levels in a sensitive population experiencing formaldehyde stress (exposed to 4 mM formaldehyde for 4 hours). Log_2_ fold change is the mean of 3 biological replicates. Full data for all genes in the *M. extorquens* genome, and standard error estimates, are given in the Supporting Information, Data S1.

In organisms where mechanisms of formaldehyde stress response have been characterized, formaldehyde resistance is achieved by formaldehyde detoxification via direct oxidation to formate or by pathways that are dependent on thiols, pterin cofactors, or sugar phosphates [29, 42]. Here, we found no significant changes in the regulation of any C_1_ metabolism genes or genes that might otherwise contribute to formaldehyde detoxification (eg, glutathione biosynthesis, alcohol/aldehyde dehydrogenases). These findings are consistent with the persistence of high concentrations of formaldehyde in the growth medium (Fig. 2) and further support the conclusion that the mechanism of formaldehyde resistance is not formaldehyde consumption.

For most of the 23 genes differentially expressed in the tolerant subpopulation, the association with formaldehyde tolerance is not immediately apparent from their annotations. However, we did find upregulation of four genes encoding UspA (universal stress protein) domain proteins and one Hsp20 (heat shock protein) gene (Table 1). Additionally, a three-gene operon encoding cytochrome *bd* ubiquinol oxidase was upregulated; this complex is classically associated with respiratory electron transport but has more recently been shown to contribute to bacterial tolerance of several stresses such as high temperature, oxidative stress, and cyanide stress [43, 44]. The 15 remaining differentially expressed genes encode transport-associated proteins (4), regulatory proteins (3), signaling proteins (3), hypothetical proteins (4), or proteins with domains of unknown function (1). Notably absent are genes involved in DNA repair, as genotoxicity is often presumed to be the mechanism of formaldehyde-induced cell death [45] and DNA repair systems are upregulated in other bacteria upon formaldehyde exposure [42]. Collectively, our results indicate that the formaldehyde tolerance phenotype in *Methylobacterium* may involve stress response mechanisms that are similar to those observed in other organisms for other stressors and may involve the coordination of several cellular processes.

### Tolerant subpopulation behavior is observable at the single-cell level

As a final step in confirming single-cell phenotypic heterogeneity in formaldehyde tolerance, we observed growth of single cells directly using high-resolution time-lapse phase-contrast microscopy of cultures embedded in agar pads. This method allowed us to observe the division times and potential morphological aberrations of individual cells and their progeny in microcolonies over the first 12 hours of growth. Because the cells tolerant to 4 mM formaldehyde are only present at a frequency of approximately 10^-4^ in the initial population, we conducted formaldehyde exposure at a lower concentration, 2.5 mM formaldehyde, at which plating experiments (see below) suggested that ∼1-4% of cells would be able to grow. Indeed, we found that 11 out of 546 cells (1.97%) were able to grow at that formaldehyde concentration, compared to 100% of the 256 cells observed in the no-formaldehyde condition (Fig. 6). In addition, all of the cells that grew in the presence of formaldehyde did so normally: we found no significant effect of formaldehyde on cell division time (median doubling time at 0 mM: 2.58 hours; at 2.5 mM: 2.58 hours; *p*=0.262 by Mann-Whitney *U*-test). We did observe that cells in formaldehyde took slightly longer to complete the first division (median lag time was 1.25 hours later in formaldehyde; *p*<0.001, Mann-Whitney *U*); this may indicate minor cell damage or a modest inhibitory effect of formaldehyde in the medium, or it may have resulted from slight differences in the preparation of the cells (as it was technically infeasible to conduct the two experiments simultaneously). Of the variance in doubling times among individual cells, most was explained by the microcolony to which the cell belonged (*p*=0.001, PERMANOVA) but not by formaldehyde treatment (*p*=0.323), indicating that there is some heritability in growth rates (Fig. 6). Finally, we did not witness any partial-growth or impaired-growth phenotypes: the 535 cells that were unable to grow in the presence of formaldehyde showed no detectable elongation or other change in morphology.

**Fig 6.**
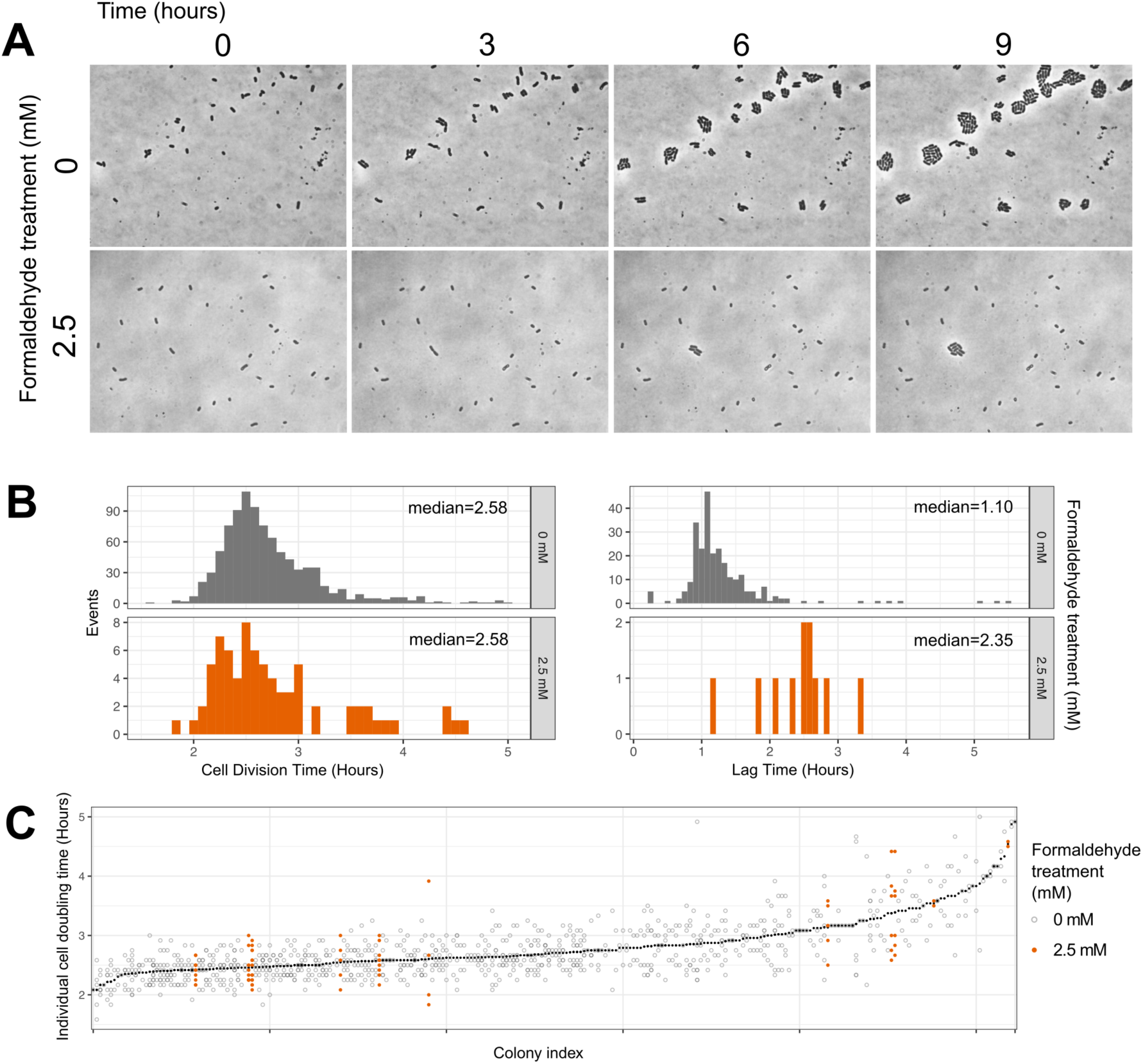
Time-lapse microscopy reveals binary (i.e., growth or non-growth) phenotypes in response to formaldehyde. A) Example images: cells were embedded in agar medium with methanol and either 0 mM (top) or 2.5 mM (bottom) formaldehyde and monitored for 9 hours (∼3 generations). At 0 mM, 256 cells were observed and all underwent at least one doubling; at 2.5 mM, 546 cells were observed and 11 (1.97%) underwent at least one doubling, in accordance with our predictions for this formaldehyde concentration (see Fig. 7). B) Histograms of cell division time (across all generations) and lag time (time between deposition and first cell division, for each microcolony) for cells that grew. No difference was observed in cell division time between the two treatments (*p*=0.262, Mann-Whitney Wilcoxon test). However, cells in formaldehyde took approximately 1.25 hours longer to reach the first cell division (*p*<0.001, Mann-Whitney). C) Scatterplot of individual cell doubling times; each position along the x-axis represents a single microcolony, ordered by mean doubling time (shown in black symbols). Individual doubling time of each cell was strongly predicted by the colony it came from (*p*=0.001) but not by formaldehyde treatment (*p*=0.323, PERMANOVA).

### Substrate affects the growth rate of tolerant cells

Our single-cell observations of growth suggested that formaldehyde-tolerant cells grow normally; the formaldehyde tolerance phenotype thus differs from other microbial stress-tolerance phenotypes previously described (e.g., sporulation, persistence) in that it does not require cells to sacrifice proliferative ability by entering a non-growing state in order to survive stressful conditions. Moreover, our observations of colony appearance times suggested that tolerant cells avoid the damage incurred by formaldehyde-sensitive cells. We therefore investigated the possibility that other fitness tradeoffs might exist, by comparing non-selected *M. extorquens* cells with those of a tolerant population (i.e., selected by growth in liquid culture at 4 mM formaldehyde).

We observed no difference in the resistance of the tolerant population to several other chemical stressors (hydrogen peroxide, and the antimicrobial compounds rifampicin, vancomycin, cefoxitin, novobiocin, nalidixic acid, ciprofloxacin, erythromycin, kanamycin, gentamicin, chloramphenicol, colistin) during growth on agar medium (data not shown). However, we did observe differences in growth rate when comparing the tolerant and naive (non-selected) populations side-by-side in liquid batch culture, depending on whether the growth substrate was methanol (15 mM) or the non-methylotrophic substrate succinate (3.5 mM). In methanol medium, the two populations grew at similar rates (naive: *r*=0.222±0.023; tolerant: *r*=0.202±0.051 h^-1^); however, on succinate medium, the naive population grew faster (naive: *r*=0.270±0.002 h^-1^; tolerant: *r*=0.215±0.030 h^-1^) (Fig. S3). The differences in growth rate among the four treatments were in this case not found to exhibit statistically significant differences by ANOVA (*F*=2.617, *p*=0.123 for the model; *p*=0.940 for the planned contrast between the two populations on methanol and *p*=0.136 on succinate). However, the apparent slight advantage of the naive population over the tolerant on succinate was attributable to the fact that the naive population grew faster on succinate than on methanol, consistent with observations commonly made in our lab (Fig. S7) and reported in the literature [28]. If the formaldehyde-tolerant population does not show the typical increased growth rate on succinate, it could indicate that tolerance is associated with methylotrophic metabolism, and may provide a clue as to the conditions in which tolerant cells could be out-competed by sensitive cells in the environment.

### Formaldehyde tolerance is a continuous phenotype

As described previously, many cases of phenotypic heterogeneity display binary phenotypes (e.g., persistent or not [13]). On the other hand, a few display a continuous distribution (e.g., resistance to chloramphenicol along a gradient of possible MBC levels [24]). While formaldehyde exposure results in a binary outcome for each cell—either growth, or cessation and death—we sought to quantify the ratio of these outcomes along a gradient of formaldehyde concentrations, by plating *M. extorquens* onto agar medium containing formaldehyde at concentrations between 1 and 10 mM in increments of 1 mM. The results showed no evidence of a bimodal distribution along the concentration axis; rather, tolerance is continuous, peaks at 0 mM, and declines exponentially and predictably with increasing formaldehyde concentration (Fig. 7). We were able to detect cells with tolerance levels as high as 6 mM. While we found growth stage to have an effect on the abundance of tolerant cells (populations in exponential growth were shifted toward slightly higher tolerance), the qualitative shape of the distribution remained the same. Furthermore, populations that were cultured in the same way and sampled at the same growth stage reproduced similar formaldehyde tolerance distributions across multiple days (Fig. S4). Notably, all colonies that grew on formaldehyde agar medium were of uniform shape and size, suggesting that tolerant cells shared similar appearance times and growth rates, as was previously observed at 4 mM (Fig. 4). In addition, cell proliferation assays with the membrane-intercalating dye performed at concentrations of 2, 3, and 5 mM formaldehyde qualitatively recapitulated the previously described population turnover seen with the assay at 4 mM, and supported the relationship between concentration and abundance of tolerant cells (Fig. S5). Thus, although the consequence of formaldehyde upon an individual cell is binary (growth or not, rather than a range of growth rates), the distribution of tolerance (i.e., MBC values) within a population is continuous.

**Fig 7.**
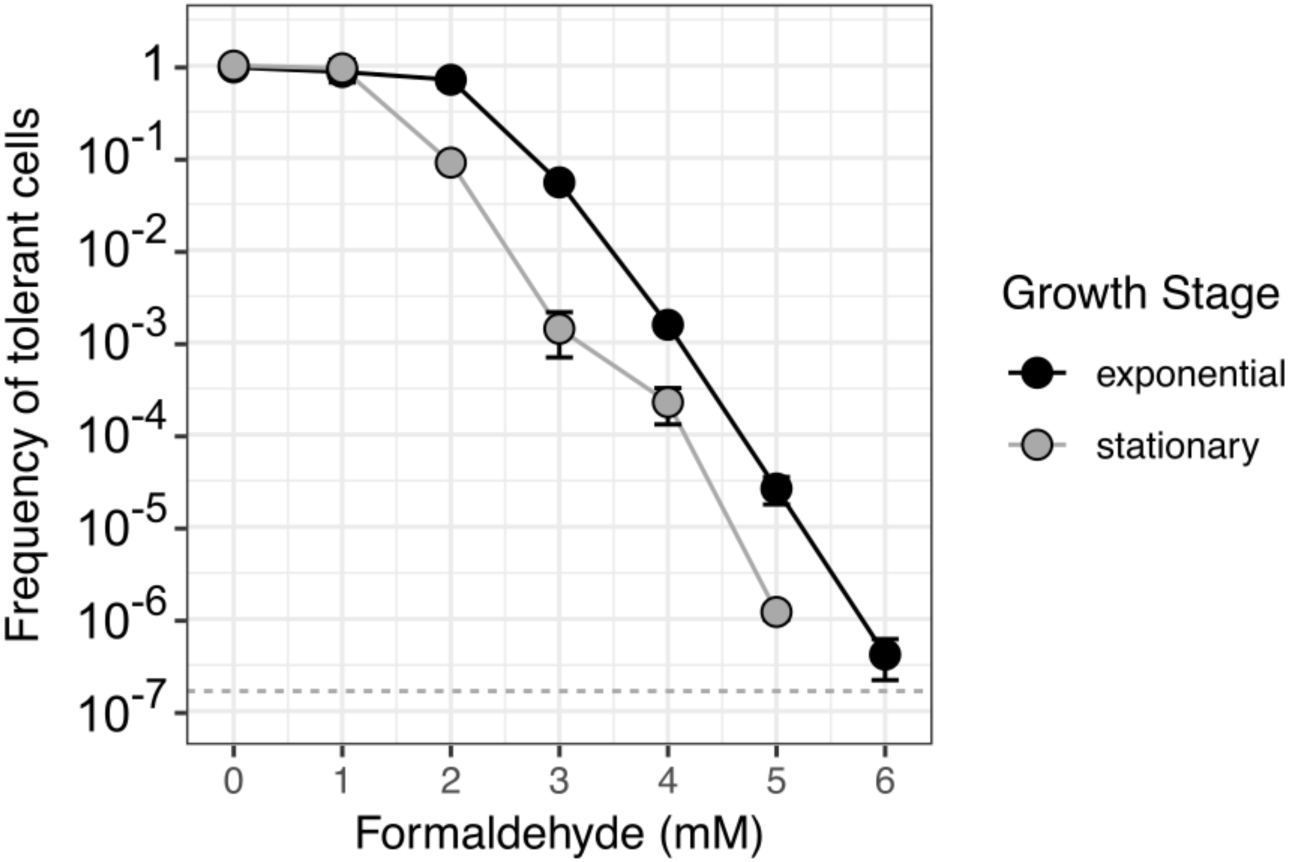
Subpopulations of formaldehyde-tolerant cells are distributed within a wild-type population with continuous, exponentially-decreasing frequency. *M. extorquens* cells not previously exposed to formaldehyde were plated onto methanol agar medium containing a range of formaldehyde concentrations at 1-mM intervals. The frequency of tolerant cells is expressed as the ratio of the colony-forming units (CFU) on formaldehyde medium at the specified concentration to the CFU on formaldehyde-free (0 mM) medium. Error bars denote the standard deviation of replicate experiments from 5 different dates (shown individually in Fig. S4). Detection limit is indicated by the dashed horizontal line.

### Transitions in tolerance phenotypes over time depend on growth conditions

Given our previous observations that growth conditions can change the shape of the formaldehyde tolerance distribution in a population, and that formaldehyde tolerance, once selected for, can be lost, we sought to characterize more precisely the processes by which tolerance distributions might shift in a population. Specifically, we hoped to better understand the relative importance of formaldehyde-mediated selection (whereby formaldehyde exposure kills sensitive cells) and active changes made by cells to alter phenotype. We began by monitoring the abundance of each subpopulation, at tolerance levels between 0 and 10 mM in 1 mM intervals, over time during a 4 mM formaldehyde exposure experiment. We monitored for 20 hours, the time it takes for most sensitive subpopulations to lose viability entirely (Fig. 2). As expected, we found that in these conditions, all subpopulations tolerant to <4 mM decrease in abundance and the subpopulations tolerant to ≥4 mM increase, resulting after 20 hours in a tolerance distribution with a maximum at 4 mM (Fig. 8, S6).

**Fig 8.**
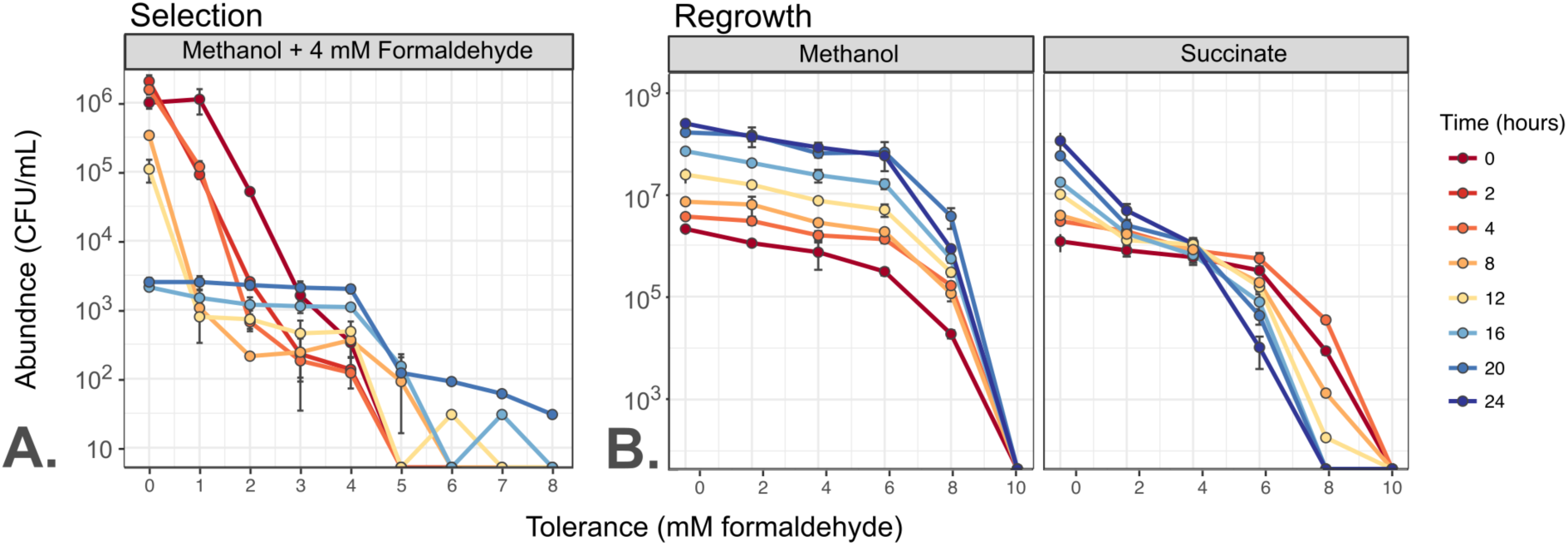
The distribution of formaldehyde tolerance within an *M. extorquens* population changes over time depending on growth conditions. Plots show total abundance (not frequency) of cells tolerant to each level of formaldehyde, as assessed by plating onto selective medium; each colored line represents one timepoint and error bars represent the standard deviation of three plating replicates. For clarity, only one biological replicate is shown; results from other replicates are shown in Fig. S6. Populations were tested for tolerance at up to 10 mM (Selection) or 12 mM (Regrowth), but 0 CFU were detected above 8 mM in either condition. A) Exposure of a naive population to 4 mM formaldehyde results in rapid decline of subpopulations with tolerance levels <4 mM and selective growth of subpopulations with tolerance levels ≥4 mM. B) When the population from A), enriched in tolerant cells, is transferred to medium without formaldehyde, tolerance distribution dynamics depend on the growth substrate provided. If growth occurs on methanol, all subpopulations grow equally well: the enrichment of formaldehyde-tolerant populations is retained for the full 24 hours (∼7 generations) of observation. If growth occurs on succinate, subpopulations with high tolerance decline in abundance and those with low tolerance increase: the population reverts to its original naive distribution.

In order to measure the rate at which phenotypic tolerance in the population returns to its original distribution, we conducted a formaldehyde-free regrowth experiment. Specifically, we transferred the selected, high-tolerance population to liquid medium without formaldehyde and monitored the changes in tolerance distributions for the next 24 hours in two different conditions: one with methanol as the sole carbon substrate, and the other with succinate. We observed a marked difference between the two growth conditions in their effect on population tolerance distributions over time. In the succinate medium, only the populations with low tolerance increased in abundance, whereas those with high tolerance *decreased* in abundance, so the shape of the distribution shifted back toward that of naive *M. extorquens* cells (Fig. 8, S6). The observation that tolerant cells decreased in abundance even during an increase in the overall population suggests that cells were shifting in phenotype from high tolerance to low. In contrast, in the methanol medium, all tolerant subpopulations increased in abundance at the same rate: the overall shape of the distribution, with its high proportion of tolerant cells, stayed the same. Growth in methanol medium thus maintains phenotypic formaldehyde tolerance in a population that is *already* tolerant, even though it does not induce or select for tolerance in sensitive populations. This unexpected substrate-based hysteresis (historical dependence) may be due to the small amount of formaldehyde produced inside the cell during methylotrophic metabolism, which might trigger cells, either through a stress-response mechanism or through regulation of methylotrophic metabolism, to remain in a tolerant phenotype even if external formaldehyde is not present in the growth medium.

### Mathematical modeling elucidates cellular phenotype transition processes

To better understand what biological processes might be responsible for the observations of the three experiments described above, we developed a mathematical model to test several hypotheses. We examined the dependence of death rate upon formaldehyde concentration in the medium and tolerance phenotype of the cell. We also asked whether shifts in the tolerance distribution could be explained by growth and selective death alone, or involved other processes. To this end, we tested the effect of introducing two processes by which cells might actively transition along the 1-dimensional axis of “phenotypic space”: one involving random phenotype transitions in any direction (a Brownian motion, or diffusion, process), and one involving directed transitions toward either higher or lower tolerance (an advective process) (Fig. 9) (An R notebook containing all model code is given in Supporting Information File S1, and data in Data S2 and Data S3). We evaluated models based on their ability to reproduce the dynamics from the three experimental conditions shown in Figure 7: selection during a 4 mM formaldehyde exposure experiment, and formaldehyde-free regrowth of the selected high-tolerance population on either succinate or methanol.

**Fig 9.**
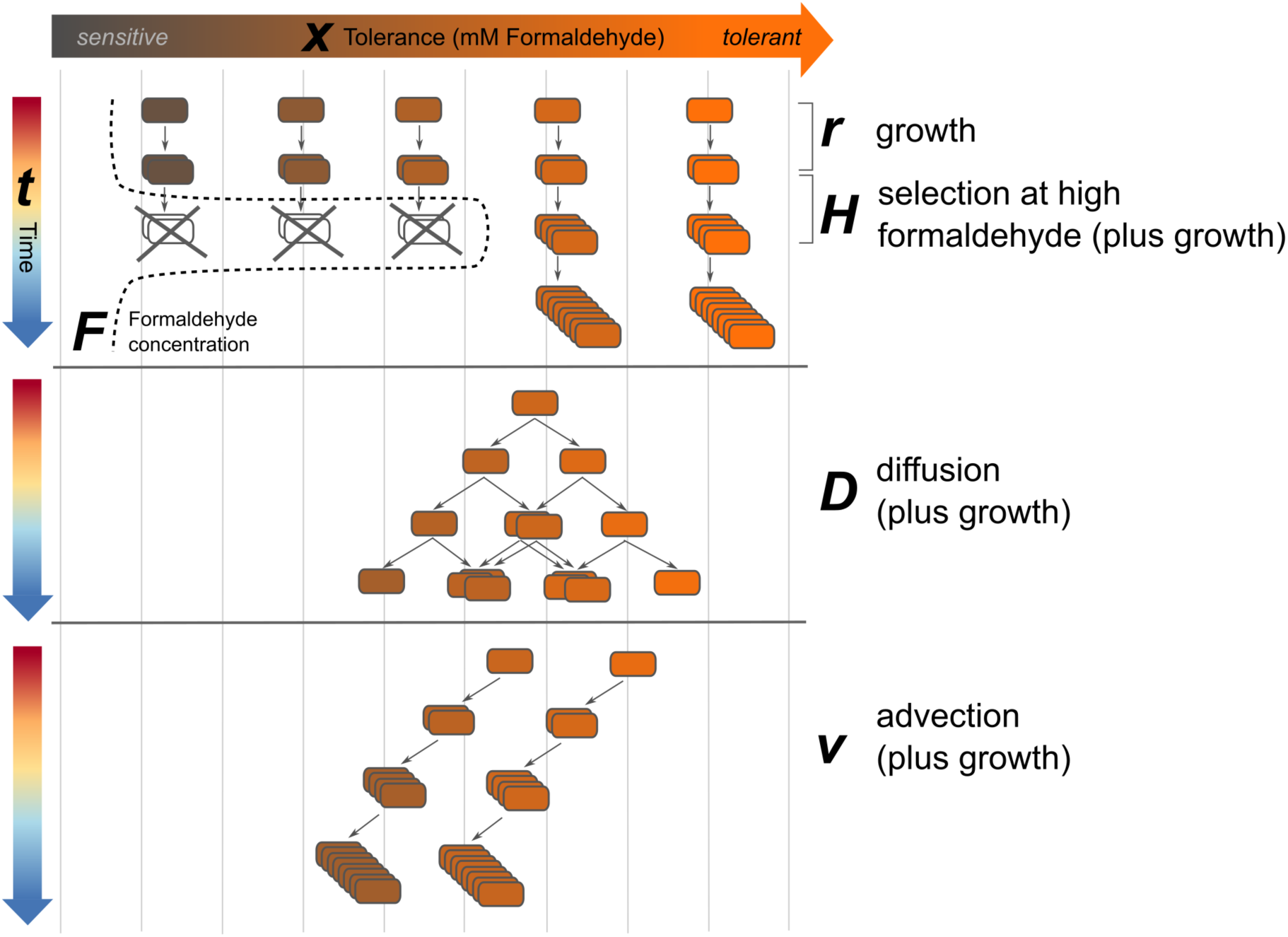
Schematic of processes described by mathematical model of tolerance distribution dynamics. Cells exist in 1-dimensional phenotype space along a continuum from sensitive to tolerant, with *x* denoting the maximum concentration of formaldehyde (*F*) at which a cell can grow. Under normal growth (at rate *r*), progeny cells carry the same tolerance phenotype as their parents. Exposure to formaldehyde results in the death of low-tolerance (*x<F*) phenotypes at a rate described by *H(x, F).* In the process of diffusion, cells and their progeny shift to adjacent tolerant states according to the diffusion constant *D*, resulting in the broadening of the population’s tolerance distribution. In advection, cells and their progeny move in a single direction in tolerance space at rate *ν*, resulting in an overall shift in the population’s distribution toward either lower or higher average tolerance.

We modeled the dynamics of *M. extorquens* populations during exposure to formaldehyde with a partial differential equation:

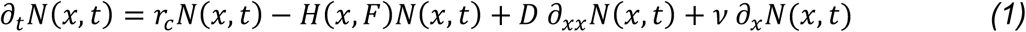

The population is structured by phenotype, with *N*(*x,t*) denoting the concentration of cells (CFU*mL^-1^) with formaldehyde tolerance *x* (mM) at time *t* (hours). The formaldehyde tolerance of a cell is defined as the maximum concentration of formaldehyde in which the cell can grow. The model tracks cells in a well-mixed, closed population as they grow on substrate *c* at per capita rate *r_c_* (h^-1^), die at per capita rate *H*(*x,F*), and change phenotype (Fig. 9). We assume that phenotypic transitions potentially occur due to two processes, diffusion with coefficient *D* (mM^2^*h^-1^) and advection with rate *v* (mM*h^-1^) (where mM refers to tolerance level). The death function *H(x,F)* describes per capita death rate of cells as a function of formaldehyde concentration and is given by:

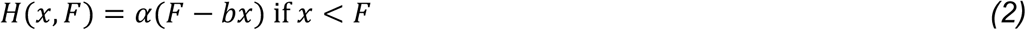

where *F* (mM) is the formaldehyde concentration, *α* (mM^-1^*h^-1^, where mM refers to formaldehyde in the medium) is the death rate, and *b* (mM tolerance / mM formaldehyde) specifies the sensitivity of the death rate to a cell’s formaldehyde tolerance level.

For each of the three experimental conditions separately, we used maximum likelihood to fit the parameters *α*, *b*, *D*, and *v* to the data, and used a likelihood ratio test on the nested models to determine the best model structure. For the selection scenario, we began with a 1-parameter model with *α* > 0 and the other parameters equal to 0, and tested 2-, 3-, and 4-parameter models sequentially, at each step choosing the model with the highest likelihood as long as it was significantly better than the simpler model (results in Table 2). For regrowth (where there is no death), we fit only *v* and *D*. In each of the three experimental conditions, our model was able to reproduce the experimental observations extremely well (pseudo-*R*^2^ = 0.973, 0.993, 0.991 for the formaldehyde selection, methanol regrowth, and succinate regrowth conditions respectively) (Table 2).

**Table 2.** Comparison of possible models describing formaldehyde-tolerance phenotype transition processes of M. extorquens populations. For each combination of culture condition and growth substrate, we used a stepwise procedure to evaluate nested models using a likelihood ratio test. Shown below are the best-fit values for the four fitted parameters in each of those models, and test results for each model. *α*: dependence of death rate on formaldehyde (h^-1^*mM^-1^). *v*: advection rate (mM*h^-1^). *D*: diffusion constant (mM^2^*h^-1^). (for *v* and *D*, mM denotes tolerance). *b*: dependence of death rate on individual tolerance level (mM tolerance / mM formaldehyde in medium). For the likelihood ratio test, name of the model used as the null model, as well as the χ^2^ value and *p*-value, are given. Gray shading: the best-supported model for that experimental scenario. Pseudo-R^2^ values for those models were: for formaldehyde selection, 0.973; for methanol regrowth, 0.993; for succinate regrowth, 0.991.

| Model | Experimental scenario |  | Parameters |  |  |  | Likelihood Ratio Test |  |  |
| --- | --- | --- | --- | --- | --- | --- | --- | --- | --- |
| | Condition | Substrate | $\alpha$ | $b$ | $v$ | $D$ | null model | $\chi^2$ | $p$ |
| F1 | selection | Methanol + Formaldehyde | 0.152 | n/a | n/a | n/a | n/a | n/a | n/a |
| F2a | selection | Methanol + Formaldehyde | 0.177 | 0.800 | n/a | n/a | F1 | 0.914 | 0.339 |
| F2b | selection | Methanol + Formaldehyde | 0.156 | n/a | -0.020 | n/a | F1 | 17.455 | 3E-05 |
| F2c | selection | Methanol + Formaldehyde | 0.160 | n/a | n/a | 0.026 | F1 | 18.480 | 2E-05 |
| F3a | selection | Methanol + Formaldehyde | 0.202±0.022 | 0.770±0.147 | n/a | 0.019±0.006 | F2b | 9.379 | 0.002 |
| F3b | selection | Methanol + Formaldehyde | 0.165 | n/a | -0.034 | 0.007 | F2b | 1.647 | 0.199 |
| F4 | selection | Methanol + Formaldehyde | 0.204 | 0.791 | 0.007 | 0.022 | F3a | 0.029 | 0.865 |
| M0 | regrowth | Methanol | n/a | n/a | n/a | n/a | n/a | n/a | n/a |
| M1a | regrowth | Methanol | n/a | n/a | 0.018±0.006 | n/a | M0 | 7.843 | 0.005 |
| M1b | regrowth | Methanol | n/a | n/a | n/a | 4.94x10<sup>9</sup> | M0 | 0.000 | 1.000 |
| M2 | regrowth | Methanol | n/a | n/a | 1.85x10<sup>-2</sup> | 4.60x10<sup>-8</sup> | M1a | -0.008 | 1.000 |
| S0 | regrowth | Succinate | n/a | n/a | n/a | n/a | n/a | n/a | n/a |
| S1a | regrowth | Succinate | n/a | n/a | 0.189 | n/a | S0 | 97.197 | <0.001 |
| S1b | regrowth | Succinate | n/a | n/a | n/a | 6.43x10<sup>9</sup> | S0 | 0.000 | 1.000 |
| S2 | regrowth | Succinate | n/a | n/a | 0.285±0.026 | 0.033±0.010 | S1a | 15.918 | <0.001 |

### Phenotype transition processes change according to growth conditions

We examined not only what the best-fit value was for each of the four parameters of interest, but also whether there was support (by likelihood ratio test) for including each of the parameters. The three experimental conditions differed from one another in both considerations, suggesting that the rate and nature of phenotype transition processes change depending on environment.

In the formaldehyde selection regime, the best model included both death parameters (*α*=0.202±0.022 h^-1^, *b*=0.770±0.147), but for the phenotype transition parameters, only diffusion (*D*=0.019±0.006 mM^2^*h^-1^) and not advection. This indicates that the changes we observed in the formaldehyde tolerance distribution during formaldehyde exposure are due not only to death, but also involve phenotype shifts consistent with diffusion. Diffusion leads to the spread of phenotypes consistent with what was observed in the model after 20 hours, including the presence of population density at *x*=2 and *x*=3 mM (which otherwise would have decreased to below detection in the absence of transitions from higher-tolerance phenotypes) and some cells at *x*>6 (which were not observed in the initial population, but may have resulted from transitions from lower-tolerance phenotypes) (Fig. 10).

**Fig 10.**
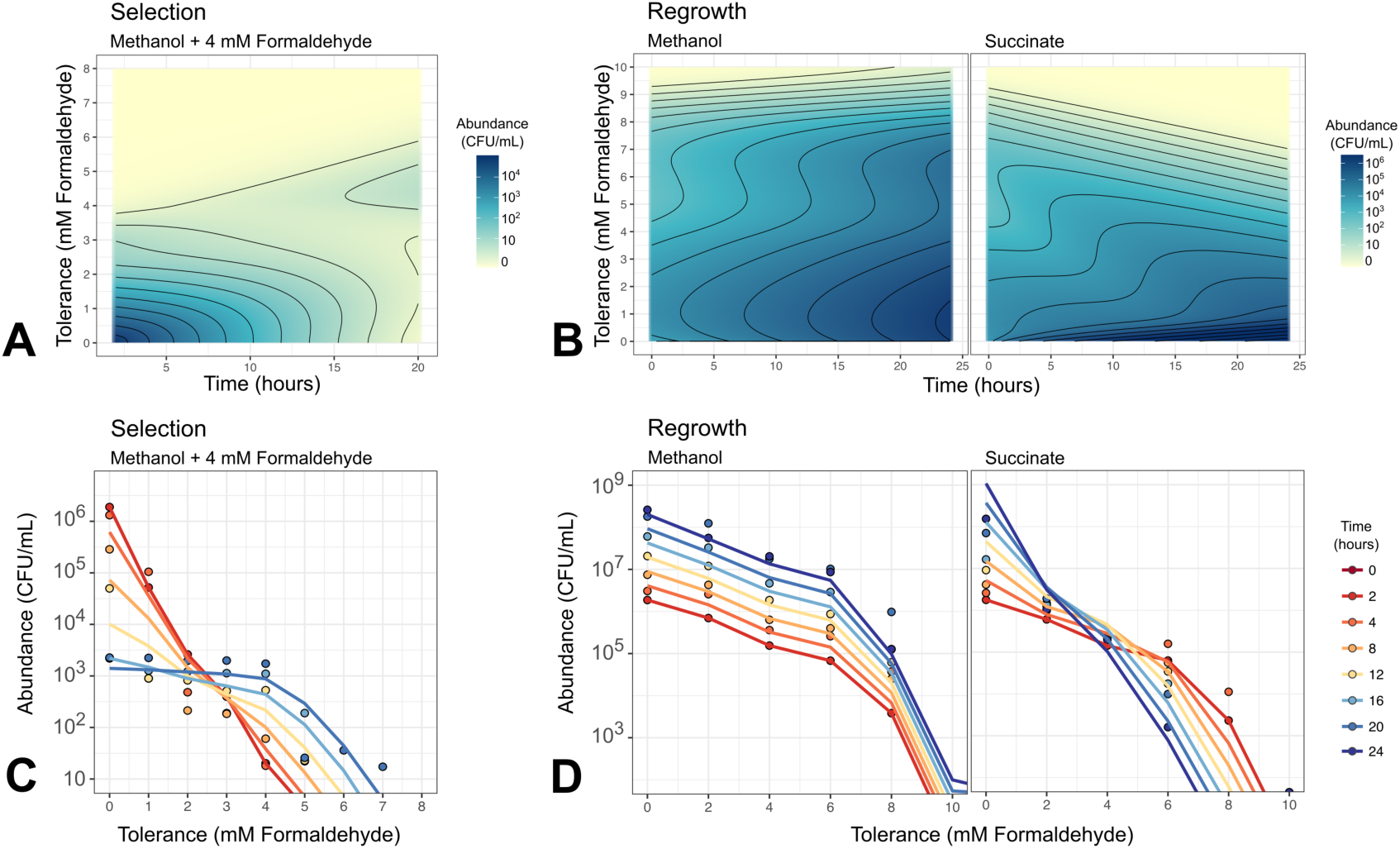
Mathematical modeling reproduces growth, death, and phenotype transition dynamics of *M. extorquens* population under multiple conditions. A) and B) Heat maps showing model simulations of population dynamics. Model parameters as given in Table 2. Note that model results are continuous in phenotype space, and non-cumulative (the abundance at concentration *x* shows only the number of cells for which that is the maximum concentration tolerable, not the number of all cells that can grow at that concentration; see Methods for details). C) and D) Comparison of model results (lines) and experimental data (points). Experimental data are averages of 3 biological replicates; model results have been binned at 1- or 2-mM intervals, and summed to form cumulative distributions, to facilitate comparison. A and C) 4 mM formaldehyde exposure experiment, resulting in selection of cells with >4 mM tolerance. B and D) Formaldehyde-free regrowth experiment, in which the selected high-tolerance population is transferred to medium without formaldehyde and either methanol or succinate as the carbon substrate, resulting in different shifts in phenotype distribution.

For the scenarios involving regrowth of a selected population on formaldehyde-free medium, the parameter estimates were markedly different from those in the selection condition, and from each other. For methanol growth, we found support only for very mild advection toward lower tolerance (*v*=0.018±0.006 mM*h^-1^). For succinate growth, we found support for both advection (*v=*0.285±0.026 mM*h^-1^) and diffusion (*D*=0.033±0.010 mM^2^* h^-1^); the advection term for succinate was an order of magnitude greater than that for methanol, indicating strong shifts in the direction of lower tolerance when high-tolerance cells are grown on succinate. This agrees with our earlier qualitative observations that growth on succinate, but not on methanol, leads *M. extorquens* populations to lose formaldehyde tolerance rapidly and in general to undergo diversifying phenotype transitions. Furthermore, it supports our hypothesis that formaldehyde tolerance is associated with methylotrophic growth, and implies that further work is needed not only to understand the mechanism of phenotypic formaldehyde tolerance shifts in *M. extorquens* but also the regulation of those processes.

The inclusion of *b* in the model specifically allowed us to test the possibility that a cell’s formaldehyde tolerance might determine not only the threshold concentration above which it dies, but also the rate at which it dies (Fig S8). That is, if *b*=0, the death rate is equivalent for all cells regardless of tolerance level as long as tolerance is below the threshold; but for 0<*b*≤1, cells with lower tolerance levels die more quickly than those with higher tolerance, and the strength of this dependence upon *x* scales with the value of *b*. The value of *b* that best fit our data was 0.770 (Table 2), indicating that formaldehyde-tolerant cells may receive some protection from the tolerance phenotype even at concentrations above their MBC. However, both the experimental results and the model simulation showed a bimodal phenotype distribution after formaldehyde exposure, consistent with a weak dependence of death rate from tolerance level (Fig. S8). Future experiments with a greater number of observations at low-tolerance phenotypes would better elucidate the relationship between tolerance and death dynamics.

## Discussion

We have described here a novel example of phenotypic tolerance to a metabolic toxin that is continuously distributed across individual cells in a clonal microbial population, where the phenotypic heterogeneity is present even when all cells inhabit the same environment, but its distribution undergoes dynamic shifts when the population experiences different environmental conditions. Wild-type populations of genetically identical *Methylobacterium extorquens* cells grown in well-mixed liquid medium contain individuals with maximum formaldehyde tolerance levels ranging from 0 mM to 6 mM (and tolerance at up to 8 mM has been observed after selection); although the distribution is continuous and exponentially decreasing, individuals show binary growth/non-growth phenotypes at any given formaldehyde concentration, and may transition among phenotypic states through both bidirectional phenotype diversification and responsive, directed phenotype shifts.

When might *M. extorquens* experience formaldehyde in its natural environment, and at what concentrations? *M. extorquens* excretes formaldehyde during the first stages of the switch between multi-carbon and single-carbon metabolism [27]; in addition, formaldehyde is a metabolic intermediate in the consumption of many lignin-derived aromatic compounds [46, 47], and we have observed lignin degraders to excrete formaldehyde into the growth medium at millimolar levels during growth in batch liquid culture on methoxylated aromatic compounds [48]. Thus, it is possible that, in the environment, formaldehyde concentrations in the millimolar range might accumulate transiently on the microscale, especially within cell aggregates such as those observed on plant leaves [49]. However, it is also possible that formaldehyde tolerance is an outward manifestation of a cellular state that has quite a different relevance in the native environment of *M. extorquens*: high tolerance to local (external) formaldehyde may indicate a high capacity for tolerating (internal) formaldehyde generated through methylotrophic metabolism. As such, this phenomenon may provide insight into processes that are general to many organisms whose central metabolic pathways involve toxic intermediates [50, 51].

The mechanisms by which high-tolerance cells can maintain normal growth in the presence of seemingly lethal concentrations of formaldehyde, and by which that ability is transmitted to progeny, remain yet to be elucidated. Our observation of a distinct gene expression profile in the tolerant subpopulation, including the upregulation of stress response genes (universal stress proteins, small heatshock protein 20, and cytochrome *bd* oxidase) and genes with various roles (transport, regulation, signaling, hypothetical) not known to be linked to formaldehyde, suggest that phenotypic formaldehyde tolerance may involve a system of cellular stress response to damaged, misfolded proteins, and a novel mechanism of formaldehyde tolerance. However, substantial further work will be necessary to uncover the precise mechanism by which tolerant cells are able to maintain normal growth in the presence of formaldehyde, and whether this mechanism is a cause or consequence of the differences in gene expression we observed between the tolerant and sensitive cells. Quantifying gene expression in populations tolerant to a range of different formaldehyde concentrations could shed light onto the relationship between gene expression levels and the continuous nature of the tolerance phenotype distribution.

A further clue to the potential mechanism of tolerance may arise from the association between maintenance of formaldehyde tolerance and methanol growth, and the potentially lower fitness of tolerant cells during succinate growth. These observations hint at a connection between formaldehyde tolerance and methylotrophic metabolism— a connection that would not be evident from the results of gene expression analysis alone. One potential explanation for such a relationship might lie in a physiological feedback loop enabling two different outcomes from initially very small variations in the activity of C_1_ pathway enzymes, such as those of the H_4_MPT pathway of formaldehyde oxidation, which play a crucial role in removing cellular formaldehyde [30]. Previous reports have found phenotypic heterogeneity in the related strain *M. extorquens* AM1 in growth rate, cell size, gene expression levels, and ability to switch between carbon substrates [8,52,53]. Assuming that formaldehyde transport into the cytoplasm is diffusion-driven (as no means of active transport has yet been discovered), phenotypic diversity in formaldehyde oxidation capacity would result in a range of internal concentrations across cells. Furthermore, if cells that experience formaldehyde-mediated damage to proteins and other macromolecules begin to lose their capacity to oxidize formaldehyde, this would generate a positive feedback circuit at the protein level that could ultimately determine a binary outcome to whether a cell lives or dies. The fact that we observed no elongation at all from the cells that could not grow at 2.5 mM suggests that this postulated positive feedback mechanism acts on a very fast timescale (i.e., minutes, not hours). This mechanism could explain both the continuous relationship between concentration and abundance of tolerant cells, and the binary nature of the “live normally or die immediately” distribution observed at each specific formaldehyde level, similar to the topology of interactions observed in the case of chloramphenicol resistance described earlier [24].

Similar examples of bi-stable phenotypic outcomes have been described: for instance, pyrimidine-mediated colony type switching in *Bacillus subtilis* [54], and chloramphenicol resistance [24]. In these cases, the balance between the concentrations of multiple intracellular components determines the phenotype of a cell, with cells pushed away from the threshold into one of two states by a positive feedback loop, and often maintained there by hysteresis (as we observed in the maintenance of formaldehyde tolerance among cells growing on methanol, Fig. 8). Making changes that alter the ratios of the components, through control of either gene expression or of the environment, alters the frequency of cells near the threshold and thereby the ratio of cells in each of the two bi-stable states (as we observed when altering formaldehyde concentrations in the medium). Furthermore, these systems are analogous to threshold traits observed in animals, in which observed discontinuous phenotypes (e.g., two phenotypic morphs) arise from a continuously distributed underlying trait [55]. The threshold model of quantitative genetics describes a constant threshold that determines which values of the underlying trait generate each of the phenotypic morphs; evolutionary and environmental changes can lead to shifts in the underlying trait distributions that result in different ratios of the two morphs being observed in different environments. As we have shown here, the continuously distributed trait of *M. extorquens* formaldehyde tolerance similarly changes with environment; however, in this case the threshold (environmental formaldehyde concentration, which separates tolerant cells from sensitive) is external and may also be manipulated. *M. extorquens* therefore provides a convenient model system in which to probe the dynamics of threshold traits.

Both stress response proteins and metabolic enzymes are inherited during cell division, so either pathway provides a hypothesis for the heritability of formaldehyde tolerance that we observed after formaldehyde was removed. Lineage dependence has been observed in numerous cases of phenotypic heterogeneity, and is often cited as evidence that the phenotype of interest is dictated by a heritable component of the cell with a moderately long lifetime (e.g., pole age [8], stress-protective protein aggregates [56]). In these cases, the random transitions in phenotype that we modeled as diffusion can be explained by asymmetric partitioning of cell contents [57], whereas the directed transitions that we modeled as advection could be due to responsive up- or down-regulation of the production/degradation or (in)activation of the molecules involved.

The balance between stochastic and responsive phenotype differentiation processes, as mechanisms for increasing population fitness in unpredictable environments, has often been discussed as a question of the relative costs and benefits of each. Both mathematical modeling [58, 59] and laboratory evolution [19, 60] have demonstrated that either mechanism—or both simultaneously—can be selected for, with evolutionary outcomes dictated by the cost of sensing, the timing of environmental change, the reliability of environmental cues, and the spatial structure of the community. The fact that we observed non-directed general diversification in our populations in some growth conditions, and environment-responsive, directed phenotype shifts in others, strongly suggests that that differentiation in formaldehyde tolerance in *M. extorquens* is not due simply to unavoidable molecular noise, but rather it is a regulated process conferring a fitness advantage.

If it exists, the nature of such a fitness advantage remains unclear; it is tempting to ask why *M. extorquens* does not maintain a population composed solely of high-tolerance cells. The evolutionary basis for phenotypic heterogeneity in genetically identical populations of microorganisms is frequently ascribed to diversifying bet-hedging, in which a species in an unpredictably changing environment constitutively generates progeny with multiple phenotypes to ensure that at least a few will thrive in any circumstances [12, 61]. The surface of the plant leaf, the native environment of *M. extorquens*, is indeed unpredictable: cells depend for growth upon a combination of gaseous methanol excreted from plant stomata [62] and other metabolites such as simple organic acids produced by the plant or by other microorganisms [63]. The emission of methanol, dependent on the metabolic state of the plant host and the conductivity of the stomata, undergoes large temporal variation [64]. It is therefore plausible that this environment could select for heterogeneity in formaldehyde toxicity response and in growth on non-methylotrophic substrates. An alternative evolutionary explanation is division of labor, which is an advantageous strategy when a particular activity is beneficial to the population but incurs some cost to the individual carrying it out [59]. In our experiments, the tolerant subpopulation is capable of detoxifying formaldehyde from the growth medium; although in batch culture this occurs too late to prevent the sensitive cells from dying (Fig. 2), this might occur differently in a spatially-structured community such as cell aggregates on the surface of a plant leaf.

Importantly, both bet-hedging and division of labor assume fitness tradeoffs among the phenotypes being diversified. Although tolerant cells may be slower than sensitive cells at growth on a multicarbon substrate (Fig. S8), it is unclear whether this disadvantage is substantial enough to explain the low frequency of high-tolerance cells in our *M. extorquens* populations. Further mathematical modeling could be used to quantify the effects of these growth tradeoffs and to explore conditions that could lead to the evolution of the steady-state distribution that we have observed in the lab. Furthermore, while phenotypic heterogeneity has been studied in great depth both in laboratory populations and in populations simulated by mathematical modeling, there remains a dearth of research on organisms in the natural environments in which they evolved [61, 65]. Future experiments examining the dynamics of phenotypic formaldehyde tolerance among *M. extorquens* cells growing on plant leaves, and among related methylotrophs in other environmental niches, will be essential to establishing the environmental relevance of this phenomenon.

## Materials and Methods

### Bacterial strains and culture conditions

All experiments were conducted with *M. extorquens* PA1 CM2730, an otherwise wild-type strain that contains a deletion of the cellulose synthesis operon to prevent cell clumping [66]. All cultures were grown at 30 °C in MPIPES mineral medium (30 mM PIPES, 1.45 mM K_2_HPO_4_, 1.88 mM NaH_2_PO_4_, 0.5 mM MgCl_2_, 5.0 mM (NH_4_)_2_SO_4_, 0.02 mM CaCl_2_, 45.3 µM Na_3_C_6_H_5_O_7_, 1.2 µM ZnSO_4_, 1.02 µM MnCl_2_, 17.8 µM FeSO_4_, 2 µM (NH_4_)_6_Mo_7_O_24_, 1 µM CuSO_4_, 2 µM CoCl_2_, 0.338 µM Na_2_WO_4_, pH 6.7) [66], either in liquid form or with the addition of 15 g/L Bacto-Agar (BD Diagnostics) for solid medium. Growth substrates consisted either of methanol provided at 15 mM (in liquid medium) or 125 mM (in solid medium); or succinate provided at lower concentrations to ensure the same molarity of carbon as in methanol conditions: 3.5 mM succinate (in liquid) or 15 mM (in solid). Unless otherwise noted, all liquid cultures were grown as a 20 mL volume in 50 mL-capacity conical flasks, capped with Suba-Seal rubber septa (Sigma-Aldrich) to prevent the escape of volatile compounds, with shaking at 250 rpm. Extensive testing in our lab has shown that cultures grown thus are not limited by oxygen availability and grow at the same rate as in flasks with loose lids. Prior to the beginning of each experiment, cultures were streaked from freezer stock onto solid medium and allowed to grow for 4 days to form colonies, and for each biological replicate a single colony was used to inoculate 5 mL of liquid medium in a culture tube and allowed to grow 24 hours. This overnight culture, containing ∼2×10^8^ CFU/mL at stationary phase, was used as the inoculum for the growth experiment.

### Formaldehyde exposure experiments

Growth and tolerance distribution dynamics in the presence of low concentrations of formaldehyde were assessed by inoculating stationary-phase culture at a 1:64 dilution (for an initial population of ∼3×10^6^ CFU/mL) into fresh MPIPES + methanol medium containing formaldehyde at the specified concentration (4 mM unless otherwise stated). Cultures were grown under the conditions described above (20 mL MPIPES, Suba-Seal septa, shaking at 30 °C) and sampled periodically to measure cell viability and formaldehyde concentration. Sampling was conducted as follows: 100 µL of culture was removed through the septum using a sterile syringe and transferred to a trUVue low-volume cuvette (Bio-Rad) to read optical density at 600 nm using a SmartSpec Plus spectrophotometer (Bio-Rad). An additional 200 µL of culture was removed and transferred to a microcentrifuge tube. The culture was centrifuged for 1 minute at 14,000 x g; the supernatant was removed and saved for formaldehyde measurement (see below) and the cell pellet was resuspended in 200 µL fresh MPIPES medium without carbon substrate or formaldehyde. Cells were then subjected to serial 1:10 dilutions in MPIPES to a final dilution of 10^-6^. From each of the seven dilutions, three replicates of 10 µL were pipetted onto culture plate containing MPIPES-methanol solid medium to form spots (total: 21 spots per sample per plate type). In experiments measuring formaldehyde tolerance distribution, multiple plate types were used, each type containing a different concentration of formaldehyde (see “Formaldehyde tolerance distributions,” below). The spots were allowed to dry briefly in a laminar flow hood, then lids were replaced and plates were stored in plastic bags and incubated at 30 °C for 4 days before colonies were counted. For each replicate set of seven spots, the two highest-dilution spots with countable colonies were enumerated and summed, then multiplied by 1.1 times the lower of the two dilution factors to calculate the original number of colony-forming units (CFU) in the sample. For each sample, the mean and standard deviation of the three replicate spot series was calculated.

For the measurement of death rates in the presence of high concentrations of formaldehyde, cells were grown in MPIPES medium with methanol and no formaldehyde until they reached an OD_600_ of 0.1 (∼ 1×10^8^ CFU/mL, mid-exponential phase), then formaldehyde was added to the desired final concentration and immediately mixed well. Growth conditions were as described above and samples were taken every 20 minutes to measure formaldehyde concentrations and cell viability. Growth rates were measured by fitting a linear relationship between time and the binary logarithm of CFU/mL using the lm function; differences among growth rates were assessed using either the anova function or the t.test function, in the stats package.

Fresh 1 M formaldehyde stock was made weekly by combining 0.3 g paraformaldehyde powder (Sigma Aldrich), 9.95 mL ultrapure water, and 50 µL 10 N NaOH solution in a sealed tube, and immersing in a boiling water bath for 20 minutes to depolymerize. Formaldehyde was measured using the method of Nash [67]. In brief, equal volumes of sample (or standard) and Nash Reagent B (2 M ammonium acetate, 50 mM glacial acetic acid, 20 mM acetylacetone) were combined in a microcentrifuge tube and incubated for 6 minutes at 60 °C. Absorbance was read on a spectrophotometer at 412 nm. For experiments involving large numbers of samples, the same assay was conducted in a 96-well polystyrene flat-bottom culture plate (Olympus Plastics): 100 µL each of sample and Reagent B were combined in each well of a culture plate and the plate was incubated at 60 °C for 10 minutes before reading absorbance at 432 nm on a Wallac 1420 Victor2 multilabel plate reader (Perkin Elmer). A clean plate was used for each assay; each plate contained each sample in triplicate as well as a standard curve run in triplicate.

### Genome resequencing of formaldehyde-tolerant subpopulation

Whole-genome sequencing was conducted on an *M. extorquens* population selected for high formaldehyde tolerance, to determine whether there were any genetic changes associated with the formaldehyde tolerance phenotype. In addition, the genomes of two non-selected *M. extorquens* populations were prepared, sequenced, and analyzed at the same time using the same methods, as controls to enable us to distinguish mutations specific to formaldehyde tolerance from any that may have accrued in our laboratory strain since the sequencing of the published genome.

Tolerant cells were selected via a 4 mM formaldehyde exposure experiment as described above, and allowed to grow until stationary phase; the tolerance distribution of each population was assayed to confirm that cells were indeed 100% tolerant to 4 mM formaldehyde (see “Formaldehyde tolerance distributions,” below). From each of 3 replicate populations, 2 mL of culture (∼2.2×10^8^ CFU/sample) were harvested. Genomic DNA was extracted using the Epicentre Masterpure Complete DNA and RNA Purification Kit (Epicentre/Illumina), following the manufacturer protocol for DNA purification, and the three populations were pooled. Library preparation was carried out by the IBEST Genomic Resources Center at the University of Idaho (Moscow, ID): genomic DNA was used to make shotgun libraries using the TruSeq PCR-free Library Prep Kit (Illumina) with the HiSeq-length insert option (short), amplified using KAPA (Illumina), cleaned using magnetic beads (MagBio), and quantified using fluorometry prior to pooling. The pooled library was quality-checked by Fragment Analyzer (Advanced Analytical) and quantified with the KAPA Library Quantification Kit for ABI Prism (Kapa Biosystems). Sequencing was conducted in a 1×100 run on a HiSeq 4000 at the University of Oregon Genomics and Cell Characterization Core Facility (Eugene, OR); reads were demultiplexed by the facility using bcl2fastq Conversion Software (Illumina).

Genomic data was analyzed for evidence of mutations using breseq 0.32.1 [68] with default settings, using the published genome of *M. extorquens* PA1 (NC_010172 [69]) as a reference, and comparing with a similar dataset from an *M. extorquens* CM2730 population that had been grown without formaldehyde. No predicted mutations were found in the formaldehyde-tolerant population that were not also found in the non-formaldehyde population. Three genomic loci (MEXT_RS13110, MEXT_RS12285/ MEXT_RS12290 intergenic region, MEXT_RS02695) carried SNPs identified as “marginal mutation predictions” at a frequency of <33%, but further analysis revealed these to be due to assembly errors in extremely repeat-rich regions that appear in both the tolerant and control samples. Areas containing marginal mutation predictions were further checked by PCR amplification and Sanger sequencing of the original DNA. Primers used were 5’-CTCTCCGCCGAAGTGGT-3’ and 5’-GCCTTCCTCGGGTTCAAGGG-3’ (for MEXT_RS02695); 5’-CAGGGAACGCTCGTAGAGG-3’ and 5’-CCACCGTGAAACGCACCGTA-3’ (for MEXT_RS12285-RS12290); and 5’-GTAGACCGCCTCCGAGACTT-3’ and 5’-GTAGACCGCCTCCGAGACTT-3’ (for MEXT_RS13110). Genome resequencing data have been deposited in the NCBI SRA database under BioProject PRJNA504295.

### Cell proliferation assay

Growth versus non-growth phenotypes in bulk liquid culture were assessed using a cell proliferation assay with the non-toxic green fluorescent membrane linker dye PKH67 (Sigma-Aldrich). To ensure that *Methylobacterium* cells could be easily distinguished from background events in flow cytometry, experiments were conducted using *M. extorquens* CM3839, a strain identical to CM2730 but constitutively expressing the red fluorescent protein mCherry at the *hpt* locus [70]. We first conducted a 4 mM formaldehyde exposure experiment using CM3839 to ensure that its population dynamics upon formaldehyde exposure were similar to those of CM2730 (Fig. S9). For the cell proliferation assay, stationary-phase cultures were stained and washed following the manufacturer protocol for PKH67, modified only in that all centrifugation steps were carried out for 1 minute at 14,000 x g. OD_600_ was measured after staining to account for any loss of cells, and inoculation density was adjusted to ensure an initial population of ∼3×10^6^ CFU/mL. Stained cells were used in a formaldehyde exposure experiment as described above, with treatments at 0, 1, 2, 3, 4, 5, and 20 mM formaldehyde. Unstained cells were grown alongside the stained cells as a control to assess whether staining affected growth rate or viability; no measurable difference in growth was detected. Samples were taken periodically by syringe and washed of formaldehyde as described above; instead of plating for viability, they were resuspended in 1 M (3%) formaldehyde as a fixative and stored at 4 °C until analysis by flow cytometry. Immediately prior to analysis, cells were centrifuged once more and resuspended in fresh medium to remove excess formaldehyde.

Flow cytometry was conducted in the IBEST Imaging Core at the University of Idaho using a CytoFLEX S benchtop flow cytometer (Beckman Coulter); each sample was analyzed at a flow rate of 10 µL/min for 3 minutes to ensure that an equal volume was examined from each. Output was gated to allow only events with a mCherry signal (ECD-area channel, excitation: 488 nm, emission: 610/620 bp) >10^3^. Per-cell membrane fluorescence was measured in the FITC-area channel (excitation: 488 nm; emission: 525/40 bp). Data analysis was conducted in R v.3.4.3 [71] using the flowCore package [72] to interpret fcs files and the ggplot2 package [73] to generate plots.

### Formaldehyde tolerance distributions

The distribution of formaldehyde tolerance phenotypes within a population was assessed by plating cell cultures onto agar plates containing formaldehyde. MPIPES medium was prepared with agar, autoclaved, and cooled to 50 °C; then methanol and formaldehyde to the desired final concentration were rapidly mixed in, and the agar was poured into 100 mm petri dishes. The dish lids were immediately replaced and plates were cooled on the benchtop. Plates were stored at 4 °C and used within 1 week of pouring. CFU were plated and enumerated as described above. This method has a limit of detection of 1.65×10^-7^ (an abundance of 34 CFU/mL is necessary to observe 1 cell per 30 µL plated, and the total cell population tested was 2×10^8^ CFU/mL; therefore the least-abundant subpopulation that could be detected, disregarding the effects of Poisson distributions at lower λ, is one with an average frequency of 1.65×10^-7^ within the total population).

To assess whether formaldehyde concentrations in agar culture plates changed over time due to volatilization, plates containing 0, 2, 4, 6, 8, and 10 mM of formaldehyde were assayed for formaldehyde content before and after a 3-day incubation, stored together in the same bag, at 30 °C. A small amount of agar (∼0.1 g) was excised from the plate, melted, diluted 1:10 in MPIPES medium, and assayed using the Nash protocol (described above). Each plate was assayed in triplicate. No change in concentration was detected in any of the plates (Fig. S10).

### Colony appearance time and growth rates

To measure the effect of formaldehyde damage on colony time and growth rates, colony growth was monitored using Epson Perfection V600 flatbed photo scanners. Formaldehyde exposure experiments were carried out on *M. extorquens* as described above, but diluted samples were spread-plated rather than spotted onto culture plates: 100 µL of one dilution was spread on each plate. Lids were lined with sterile black felt to reduce condensation and increase contrast, and plates were placed agar-side-down on scanner beds. Each culture contained precisely 30 mL of agar medium, to ensure uniformity of nutrient supply and hydration status across all plates; temperature probes were included between plates on scanner beds to monitor temperatures for consistency. Scanners were placed in a 30 °C incubator and image acquisition was controlled by a computer running Linux Mint, using a cron job for scheduling and a custom bash script employing the utility scanimage to take images once per hour.

Images were processed using a custom Python 3.5.6 script employing scikit-image v.0.12.1 [74] to identify colonies and measure their areas in pixels. Subsequent data analysis was conducted in R: double colonies and non-colony objects were removed from the dataset, colony appearance time was measured as the first timepoint at which colony area measured greater than 100 pixels, and colony growth rate was measured by fitting a linear relationship between time and the binary logarithm of colony area using the lm function. Variability among growth rates was assessed as the median average deviation (MAD) within the colonies from a single timepoint, using the mad function in the stats package. The effect of formaldehyde exposure time on colony appearance time, colony growth rate, and MAD was calculated using simple linear models using the lm function. Further details of both image analysis and colony growth statistics are included in Supporting Information (Fig. S2).

### Gene expression analysis

Cultures were prepared by conducting a 4 mM formaldehyde exposure experiment as described above, in biological triplicate. Cultures were harvested at 4 hours of formaldehyde treatment (stressed sensitive cells); 64 hours of formaldehyde treatment or when OD_600_ reached 0.2-0.3 (tolerant growing cells); and 16 hours with no formaldehyde treatment (sensitive unstressed control). Harvesting proceeded as follows: 10^9^ cells were centrifuged in a Beckman tabletop centrifuge (specs) for 5 minutes 4,000xg at 4 °C and supernatant was removed; cells were washed in cold fresh MPIPES medium (no carbon) to remove excess formaldehyde and centrifuged once more to pellet; supernatant was removed, and the pellets were immediately frozen in liquid nitrogen. Harvest volume was 50 mL of exponential-phase culture for the two growing treatments and 150 mL of formaldehyde-stressed cells.

RNA extraction, cDNA generation, library preparation and sequencing were conducted by Genewiz. Briefly, total RNA was extracted via the RNeasy Plus Mini Kit (Qiagen); rRNA was depleted with the Ribo-Zero rRNA removal kit (Illumina); the quality of resulting RNA samples was assessed with an Agilent 2100 BioAnalyzer and Qubit assay. Once generated, the cDNA library was sequenced on an Illumina HiSeq (2×150bp) and raw data in FASTQ format was provided to investigators.

Data analysis was conducted using an analysis pipeline developed by the University of Minnesota Genomics Center and the Research Informatics Solutions (RIS) group at the University of Minnesota Supercomputing Institute. The pipeline uses FastQC [75] to assess the quality of the sequencing data, Trimmomatic [76] to remove the low quality bases and adapter sequences, HISAT2 [77] to align the curated reads to the *M. extorquens* PA1 genome [GenBank accession: CP000908.1], Cuffquant and Cufnorm from the Cufflinks package [78] to generate FPKM expression values, and featureCounts from the Rsubread R package [79] to generate raw read counts. DESeq2 was used to convert raw count data to normalized counts and all further statistical analysis were carried out using in R [80]. Differentially expressed genes were identified by comparison to the no formaldehyde treatment cells (ie, reference treatment) and had a log_2_ fold change > 1 and a False Discovery Rate (FDR) adjusted p-value < 0.001.

### Single-cell time-lapse microscopy

To analyze the formaldehyde-dependent heterogeneous response in lag-phase and elongation rates of *M. extorquens*, we employed single-cell time-lapse microscopy using a phase-contrast inverted microscope (Leica, DMi8) equipped with an automated stage. For image acquisition, we employed a 60x (PH2, NA: 0.7) magnification objective and a sCMOS camera (Hamamatsu, ORCA-Flash 4). Time-lapse imaging was performed under ambient conditions (regulated at 26 °C ± 1 °C) with a 5-minute period using automated routines (Molecular Devices, Methamorph). All reported experiments were performed once; as our setup can accommodate only one condition at a time, the 0 mM and 2.5 mM formaldehyde experiments were each conducted once on different days.

For imaging with single-cell resolution, 2 µL of 10x diluted stationary-phase cells (approximately 6×10^5^ CFU) grown in liquid MPIPES-methanol without formaldehyde were introduced on 1 mm thick agar pads pre-deposited on a glass coverslip. The culture solution was allowed to dry for approximately 10 minutes within a biosafety cabinet and was subsequently covered with a second coverslip. Individual bacteria were therefore immobilized at the agar-coverslip interface, allowing them to grow in two dimensions. Specific locations (200×150 µm^2^) were monitored in parallel, selected by evaluating the distance between individual cells at the beginning of each experiment so that expanding micro-colonies would not overlap spatially at later time points. On average, 40 microcolonies per location were imaged, with approximately 8-10 individual cells per microcolony in the final timepoint. ImageJ [81] and manual curation were employed for cell segmentation and tracking (Fig. S11).

To prepare the pads, MPIPES-agar medium was prepared as described above (“Bacterial strains and culture conditions”), and melted at 70 °C for approximately 2 hours using a convection oven. Subsequently, methanol was introduced to the agar medium to a final concentration of 125 mM. Specific to the formaldehyde tolerance experiments, formaldehyde was also added to a final concentration of 2.5 mM. In all experiments, a polydimethylsiloxane (PDMS, Sylgard 184, Dow Corning) frame was employed to cover all free edges of the agar pads. This step was employed to minimize formaldehyde evaporation during time-lapse imaging. To fabricate these membranes, PDMS monomer was mixed with its catalyst at a 10:1 ratio, degassed, and cured at 70 °C for approximately 2 hours. Subsequently, the cured PDMS was cut to an area of 25×50 mm^2^, including a 21×47 mm^2^ internal aperture, where the melted agar was subsequently introduced.

Analysis of cell doubling times and microcolony lag times was conducted in R. The wilcox.test function was used to compare the formaldehyde-treated population with the control population by Mann-Whitney Wilcoxon test [82]. The relative effects of formaldehyde treatment and colony lineage were assessed by permutational multivariate ANOVA using distance matrices (PERMANOVA) [83], with the adonis function in the vegan package [84].

For all microscopy experiments, an aliquot of the same culture used for the microscopy experiment was simultaneously grown in batch liquid culture with same concentration of formaldehyde, and CFUs in both the total and formaldehyde-tolerant populations were tracked over time, as a control. In all cases, the growth of the liquid culture proceeded as expected and the frequency of formaldehyde-tolerant cells matched that observed in the microscopy experiment.

### Assays on selected formaldehyde-tolerant subpopulations: fitness trade-offs formaldehyde-free regrowth experiments

For characterization of the formaldehyde-tolerant subpopulation, a 4 mM formaldehyde exposure experiment was conducted as described above to select for cells with a minimum tolerance level of 4 mM. Once cultures had reached stationary phase (approximately 80 hours), they were used within 4 hours for further experiments.

Assessment of tolerance to antimicrobial drugs and to hydrogen peroxide were carried out using a disk susceptibility test: liquid exponential-phase cultures grown on methanol in standard conditions were diluted to an OD_600_ of 0.3, then spread using a sterile swab onto the surface of MPIPES-methanol-agar plates and allowed to dry. A BBL Sensi-Disc (Becton-Dickinson), impregnated with one antibiotic compound at a set concentration, was then placed on top of the plate. Plates were incubated at 30 °C for 48 hours to allow a lawn to grow; the diameter of the clearing zone around the antibiotic disk was then measured and compared between the naive and selected high-tolerance populations.

To compare growth rates on different carbon substrates, three selected subpopulations, and three naive populations not previously exposed to formaldehyde, were diluted 1:64 into fresh MPIPES medium containing either methanol or succinate, and each of those was transferred into the wells of a 48-well tissue culture plate (Corning Costar) with 640 µL culture in each of three replicate wells. Plates were incubated on a Liconic LPX44 incubator shaker at 650 rpm, and OD_600_ was read using a Wallac 1420 Victor2 plate reader. Growth rates were calculated in R using timepoints during the period of exponential growth (between 5-18 hours for methanol growth and 5-22 hours for succinate), by fitting a linear relationship between time and the binary logarithm of the OD using the lm function in R. Rates were calculated for each well individually; for each biological replicate, the mean and standard deviation of the three replicate wells was calculated. Statistical tests of differences among growth rates consisted of analysis of variance using the anova function from the stats package, and planned contrasts calculated using the lsmeans and contrast functions from the lsmeans package.

To assess tolerance distribution dynamics during formaldehyde-free regrowth, three selected subpopulations were each diluted 1:64 into two batches of fresh MPIPES liquid medium, one batch containing methanol and the other succinate. Dilutions were grown in standard flask conditions as described above, but with loose caps instead of Suba Seals. Cultures were sampled every 4 hours for serial dilution and plating as described above, with each sample plated onto each of 7 plates containing different levels of formaldehyde (0, 2, 4, 6, 8, 10, or 12 mM). Plates were incubated and colonies were counted as described above.

### Mathematical model

Growth, death, and phenotype transitions of *M. extorquens* populations were modeled using a partial differential equation (PDE; Equation 1). The full code for parameter fitting and model selection is included as an R notebook in the Supporting Information, File S1. Over the time periods for which we ran the model simulation, there was no effective change in the concentration of either of the growth substrates (methanol, succinate; each of which have extremely low half-saturation constants [85, 86]) or formaldehyde (Fig. 2); these compounds were therefore not explicitly included as time-dependent variables in the model. Furthermore, given that tolerance either has no effect upon growth on methanol, or a mild effect upon succinate growth, we use a single value of *r_c_* for all values of *x*.

Modeling was conducted in R. The PDE was solved numerically by vectorized ODEs, where each ODE corresponded to a discrete bin within tolerance space of 0.01 mM formaldehyde. A finite difference grid was created using the setup.grid.1D function, and advection and diffusion were calculated using the advection.1D or tran.1D function as appropriate for the different models, in the ReacTran package v. 1.4.3.1 [87]. The vectorized ODEs were solved using the ode.1D function from the package deSolve v1.21 [88], with the lsoda method [89]. Zero-flux boundary conditions for Equation 1 are given by: *∂_x_N*(*0,t*)*=0* and *∂_x_N*(*L,t*)*=0*. The lower boundary was set as *x*=0 because tolerance cannot fall below 0. The upper boundary (*x=L*) for each growth condition was set higher than the highest experimentally observed value, sufficiently large so as not to constrain the upward transition of cells in phenotypic space.

#### Initial conditions

The experimentally measured distributions of formaldehyde tolerances in the original populations were used as initial conditions in all model runs. Because we found that populations exhibit a slight shift toward higher average tolerance within 2 hours of transitioning from the stationary (non-growing) to the exponential (growing) phase (Fig. 7), and because our model specifically focuses on growth phenomena and does not capture behavior in stationary phase, we chose the 2-hour tolerance distribution as our initial condition (Fig. S7) in order to avoid artefacts that would result from using the 0-hour stationary-phase distribution. Three biological replicates were generated and the average of the three used to generate the distribution.

Because the model is continuous, whereas the experimental data were obtained at a resolution of 1.0 or 2.0 mM formaldehyde in tolerance space, the initial conditions for the model were generated by fitting a monotone cubic spline using the function splinefun in R with method “hyman,” to interpolate cell abundances for values of *x* at intervals of 0.01 mM (Fig. S7). Subsequently, for comparison of model results with experimental results, the model results were summed in 1 mM or 2 mM intervals to re-obtain coarse bins matching the resolution of the experimental data (as in Fig. 10).

In the model, *x* denotes tolerance as the *maximum* tolerance level of a cell or subpopulation. However, our experimental CFU counts are cumulative in that regard (*i.e*., the cells that form colonies on culture plates containing 3 mM formaldehyde include those with a maximum tolerance of not just 3 mM but also 4 mM, 5 mM, and 6 mM). For the purposes of carrying out the model, we therefore transformed the empirical tolerance distributions in the experimental datasets to calculate the actual (non-cumulative) number of CFU at each phenotype level using the following formula:

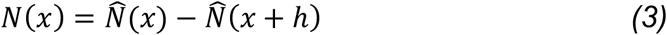

where *N*(*x*) is the number of cells that uniquely have a given tolerance level *x*, 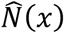 is the number of cells measured experimentally as CFU on culture plates with formaldehyde concentration *x*, and *h* is the step size between categories in data (1 mM for Fig. 8a; 2 mM for Figs. 7b-c). This transformation to the non-cumulative distribution was used after spline fitting to calculate the initial distributions for the model, and model results are shown in this format in Fig. 10a-b and Fig. S8b. However, for all other purposes, including parameter fitting as well as display in Figs. 9c-d and S12, model results were transformed back to the cumulative distribution for comparison with the experimental results. Both forms of the data are displayed in the R notebook (File S1).

A further adjustment to data from all timepoints was made to account for the experimental limit of detection (34 CFU/mL) in measuring formaldehyde tolerance distributions: growth of originally undetected low-frequency cells in the high-tolerance phenotypes could potentially be mistaken as transition into those phenotypes from lower-tolerance phenotypes. To assess whether such undetected cells would make a difference to the model results, we generated a set of “extended” CFU counts in which we made the extreme correction of adding 1 CFU to the experimental observations (the equivalent of 90.9 CFU/mL for that replicate, or 30.3 CFU/mL if averaged across the three replicates) at high formaldehyde concentrations where the observation had been 0 CFU. This correction was made to the data from all timepoints, according to the following rules:

1. If 0 colonies were observed in all technical replicates, add 1 colony to one of the replicates.
2. Only do (1) if either:

a. there is an observation of ≥1 colony at a higher concentration than the one being considered, OR
b. it is the first concentration beyond the last observation of ≥1 colony.

We carried out parameter fitting and model selection for the 4 mM formaldehyde selection experiment using both the original dataset and the extended one (as described in the previous paragraph). Although the use of the extended dataset resulted in minor differences in the estimated parameters (Tables 1 and S1, Fig. S12), the same form of the model was favored (the 3-parameter model with *α*, *d*, and *b*), and the pseudo-R^2^ (0.973) was slightly better than for the version using the original dataset (0.970). We interpreted this to mean that the inclusion or omission of rare undetected cells at high tolerance levels makes little difference to any biologically relevant conclusions, but that our correction may allow for a better model fit. We therefore proceeded with only the extended dataset for the models describing formaldehyde-free regrowth, and all parameter and statistical values given in the text are those for the extended data.

#### Parameter estimation

Growth rates (*r_c_* for substrate *c*, either methanol or succinate, h^-1^) and their standard errors were estimated from fitting experimental data of growth of a wild-type, naive population on either succinate or methanol as a primary carbon source, in the absence of formaldehyde (Fig. S7). Linear regression was used to fit the relationship between cell count and time for the exponential portion of three replicate growth curves, using the lm function in R. The growth rate on methanol was 0.195±0.001 h^-1^ and the growth rate on succinate was 0.267±0.005 h^-1^, where ± denotes 95% confidence interval.

The death rate (*α*), dependence of death on tolerance (*b*), diffusion (*D*), and advection (*ν*) parameters were estimated using maximum likelihood. Due to the exponential nature of bacterial growth, and because our data sets contain a number of zeroes, we transformed our data using the hyperbolic arcsine function 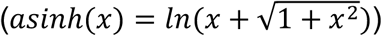 [60], which is approximately logarithmic but defined at *x*=0. The functions lm and logLik in R were used to calculate log-likelihood under a linear model; optimization was carried out in R using optim with the Nelder-Mead method. Standard errors of the parameters were calculated from the Hessian matrix. Fitted values for *D* and *v* are given in Table 2 and Table S1. In addition, we generated a phenotype-independent estimate of *α* using death rate data from a separate set of time-kill experiments (such as those in Fig. 1) at formaldehyde concentrations between 3 and 20 mM. Time-kill curves were conducted as described above, the death rate at each concentration (h^-1^) was calculated using linear regression, and the relationship between concentration and death rate was then also calculated using linear regression. By this method, we estimated *α* as 0.189±0.010 h^-^1*mM^-1^, which fell within the range of values predicted by fitting *α* for the models we tested, and within the 95% confidence interval of the estimate of *α* in the best model. This phenotype-independent estimate was not used in modeling, but helped to verify that the values we obtained by parameter fitting were reasonable.

#### Model evaluation

For each experimental condition, we used a likelihood ratio test on the nested models using a forward, stepwise procedure to choose the model that best fit the experimental data. For the formaldehyde selection scenario, the “absolute death” model (with *α* as the only parameter) was used as the null model, and each 2-parameter model (with alpha and either *b*, *D*, or *v*) was compared against it. Of the three 2-parameter models, we chose the one with the highest likelihood as long as it was significantly better than the null model. If a 2-parameter model was chosen, it became the null model and the procedure was repeated to determine whether a 3-parameter model was supported, and subsequently the 4-parameter model. For the regrowth scenarios, where no death due to formaldehyde is possible, we omitted *α* and *b*. We first compared two 1-parameter models (*v* only or *D* only) against a null model containing neither, and then evaluated whether adding the second parameter was significantly better. LR was calculated as

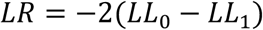

where *LL*_0_ is the log-likelihood of a reduced model and *LL*_1_ is the log-likelihood of the model being tested. Statistical significance was assessed using the chi-squared test, with degrees of freedom given by the difference in the number of parameters between the model being tested and the null model.

## Acknowledgments

We are grateful to Eric Bruger, Sergey Stolyar, Nicholas Shevalier, and Joshua Wirtz for assistance in conducting *M. extorquens* experiments; to Dipti Nayak for advice and resources facilitating those experiments; to Craig Miller for advice on analyzing colony counts; to William Harcombe and Jeremy Chacón for guidance on time-lapse imaging of colony growth using flatbed scanners; to Mark Lamourine for help with software and Dan Schneider for contributions to hardware for running the scanners; and to Pål Johnsen and Alex Bradley for helpful comments on the manuscript. Flow cytometry was conducted at the IBEST Optical Imaging Core at the University of Idaho under the guidance of Ann Norton. Genomic DNA sample preparation and sequencing were managed by the IBEST Genomics Resources Core at the University of Idaho by Dan New, Samuel Hunter, and Matt Fagnan. Juan E. Abrahante of the University of Minnesota Informatics Institute (UMII) assisted with RNA-Seq data analysis. IBEST is supported in part by NIH COBRE grant P30GM103324. Funding for this work came from an Army Research Office MURI sub-award to CJM (W911NF-12-1-0390), a CMCI Pilot Grant to CJM, CHR, and AEV (parent NIH award P20GM104420), and a grant from the John Templeton Foundation to Global Viral (Grant ID 60973). We dedicate this paper to the memory of Paul J. Joyce and his passion to bring math and biology together.

## Supporting Information

Figure S1. Formaldehyde concentrations of ≤5 mM allow growth of *M. extorquens* at a normal rate, but only after a period of lag; higher concentrations lead to longer lag times.

Figure S2. Image processing pipeline to generate colony growth data from formaldehyde-exposed cultures.

Figure S3. Formaldehyde tolerance may be associated with lower fitness on a multicarbon substrate.

Figure S4. Formaldehyde tolerance distributions in *Methylobacterium* populations are robust across experimental replicates, but vary depending on growth conditions.

Figure S5. Cell proliferation assays support the hypothesis that growth of *M. extorquens* in the presence of formaldehyde is due to a small subpopulation of tolerant cells, and that the abundance of tolerant cells decreases with increasing formaldehyde.

Figure S6. The distribution of formaldehyde tolerance within an *M. extorquens* population changes over time depending on growth conditions.

Figure S7. Estimation of growth rates and initial conditions for use in the mathematical model.

Figure S8. The parameter *b* (dependence of death rate on formaldehyde tolerance) determines the shape of the population’s phenotypic tolerance distribution after exposure to formaldehyde.

Figure S9. Cells expressing mCherry show the same formaldehyde tolerance heterogeneity as wild-type *M. extorquens* cells.

Figure S10. Formaldehyde concentrations in agar growth medium are stable over time and reflective of similar concentrations in liquid medium.

Figure S11. Time-lapse microscopy: cell segmentation and tracking.

Figure S12. Models using extended and original tolerance distributions perform similarly.

Table S1. Results of model selection using original data set for fitting (distribution not extended to account for experimental limit of detection).

File S1. Modeling phenotypic switching in *Methylobacterium extorquens*: R notebook Data S1. Excel file with original data shown in each of Figures 1, 2, 4, 5, 6, and 7.

Data S2., Data S3. Comma-separated values files with data shown in Figure 8 and used in fitting the model - required by File S1.

Data S4. Zip containing full flow cytometry results from experiment shown in Figure 3.

