## Supplemental Figures and Tables for "Microbial phenotypic heterogeneity in response to a metabolic toxin: continuous, dynamically shifting distribution of formaldehyde tolerance in *Methylobacterium extorquens* populations"

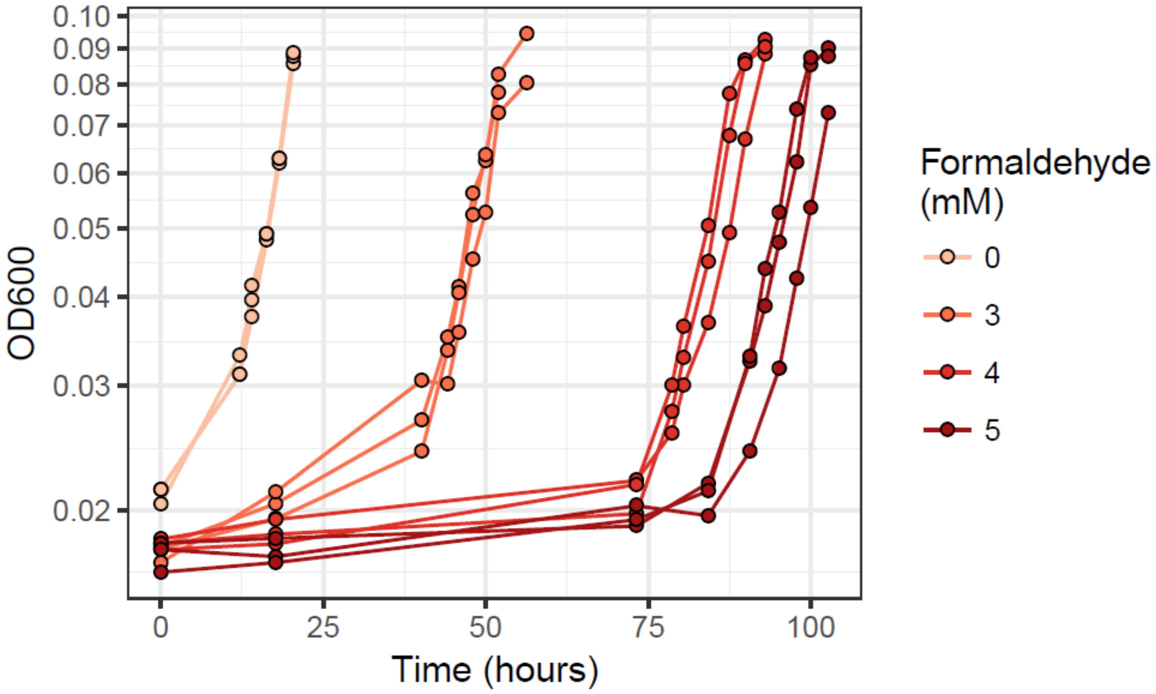


**Figure S1. Formaldehyde concentrations of ≤5 mM allow growth of *M. extorquens* at a normal rate, but only after a period of lag; higher concentrations lead to longer lag times.**
Isogenic populations of WT *M. extorquens* were inoculated into culture flasks with fresh MPIPES medium with 15 mM methanol and the indicated concentration of formaldehyde; samples were removed regularly for measurement of optical density at 600 nm. Each line represents one biological replicate.

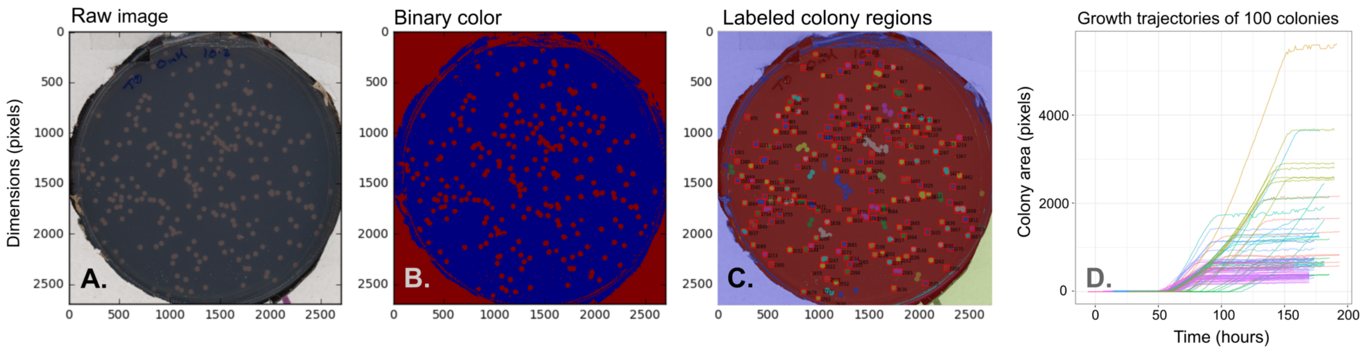


**Figure S2. Image processing pipeline to generate colony growth data from formaldehyde-exposed cultures.**

A) Color images of culture plates incubating on a flatbed photo scanner were captured once per hour; images were processed in Python 3.5.6 using a custom script employing sci-kit image v.0.12.1.

B) Each scanner bed held 6 plates; color processing was carried out for all 6 plates in an image together. The red channel only was used; the image was converted to binary color using the theshold_otsu tool to determine the threshold between light (colony) and dark (background plate) pixels, shown here as red and blue respectively.

C) For each plate, the final image was used to identify pixels belonging to each colony, using the label tool to designate colonies as individual regions. To eliminate non-colony objects and groups of multiple colonies that had grown together, a custom script was used to eliminate regions that did not meet the following criteria: 1) the region was between 40 and 10,000 pixels in size; 2) the ratio between the two dimensions of the bounding box was between 0.6 and 1.4 (approximately square), and 3) the region area occupied at least 60% of its bounding box (mostly convex). After the automatic criteria were employed, images of labeled regions were then also inspected visually, and any regions that were not single colonies were eliminated manually from downstream analysis.

D) Once the regions belonging to each colony were identified in the final image, all the images of the time series were converted to binary color and the number of bright pixels within each colony's region (the colony area) was enumerated at each timepoint. These data were imported into R v.3.4.3 for further analysis. Timepoints were adjusted to account for the fact that plates had begun growing at different times (as each population had been exposed to formaldehyde for a different length of time prior to plating). Any objects that had an area of >100 pixels at the first timepoint, or that never reached an area of 100 pixels by the final timepoint, were identified as non-colony objects and omitted from further analysis. Panel D shows growth trajectories of 100 randomly chosen colonies as an example. Colony appearance times were measured as the first timepoint at which a colony reached an area of 100 pixels or greater. Growth rates were calculated by fitting a linear relationship between time (in hours) and the binary logarithm of colony area (in pixels) using the lm function. Only growth rate estimates with an adjusted $R^2$ of >0.95 were retained.

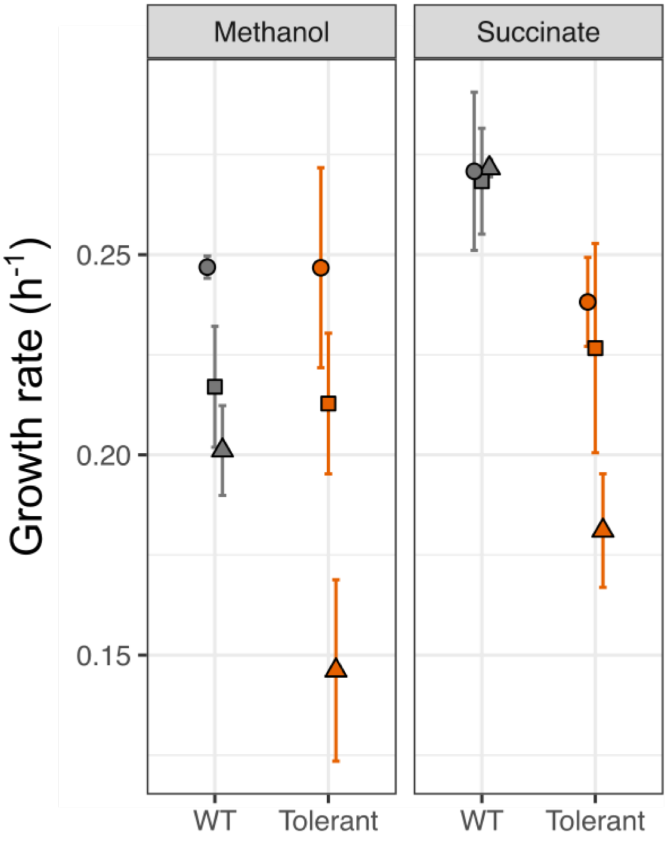


**Figure S3**. **Formaldehyde tolerance may be associated with lower fitness on a multicarbon substrate.**
Three *M. extorquens* populations with elevated tolerance were generated by selecting for tolerant cells via a 4 mM formaldehyde exposure experiment. These ("Tolerant") populations were then compared to three non-selected ("WT") populations during growth on medium without formaldehyde, with either methanol (left) or succinate (right) as the sole carbon source. Error bars represent the standard deviation of three replicate incubations of each population. Symbols represent the inocula from which the populations originated (e.g., all squares came from the same overnight culture prior to selection). On succinate only, naive *M. extorquens* populations grow marginally faster than tolerant populations, though the difference was not statistically significant (ANOVA: $F$=2.617, $p$=0.123 for the model; $p$=0.940 for the planned contrast between the two populations on methanol and $p$=0.136 on succinate).

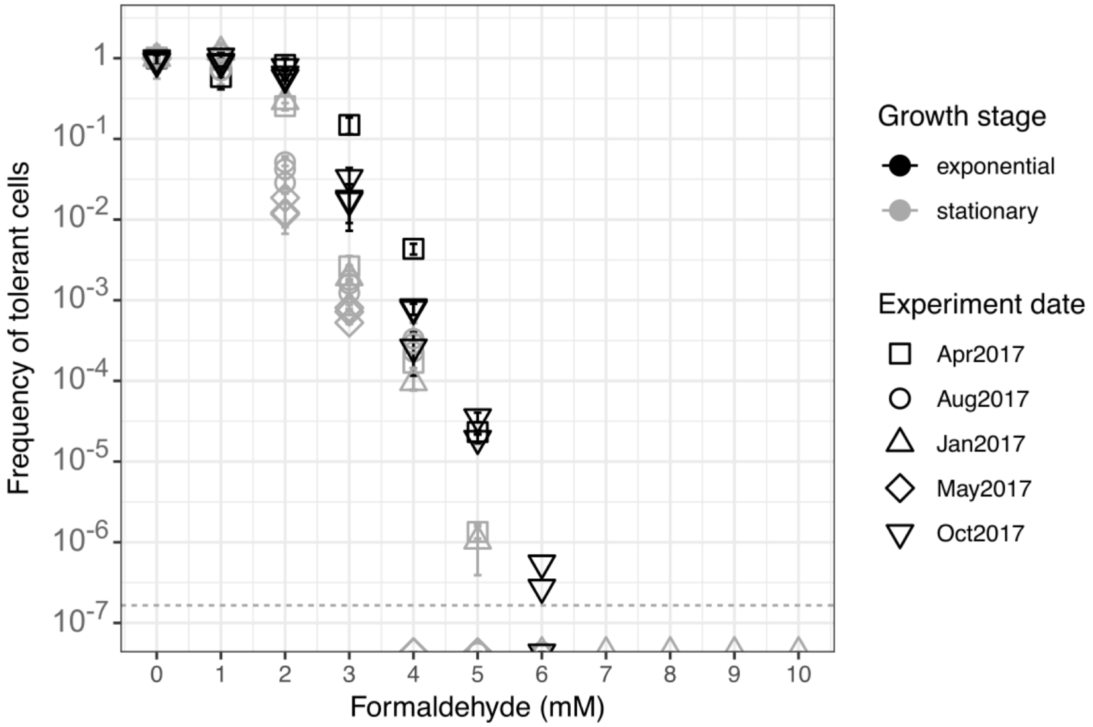


**Figure S4. Formaldehyde tolerance distributions in *Methylobacterium* populations are robust across experimental replicates, but vary depending on growth conditions.** Formaldehyde tolerance distributions were measured in isogenic *M. extorquens* populations grown on methanol medium and not previously exposed to formaldehyde, by plating onto agar methanol medium containing formaldehyde at the indicated concentrations. The frequency of tolerant cells is expressed as the ratio of the number of colonies observed on formaldehyde medium at the specified concentration to the number of colonies on formaldehyde-free (0 mM) medium. Shown here are results from several replicate experiments at different times; the average values from all replicates are shown in Fig. 6. Error bars denote the standard deviation of 3 replicate platings, and detection limit is indicated by the dashed horizontal line. In general, stationary-phase populations have lower overall tolerance than exponential-phase (actively growing) populations.

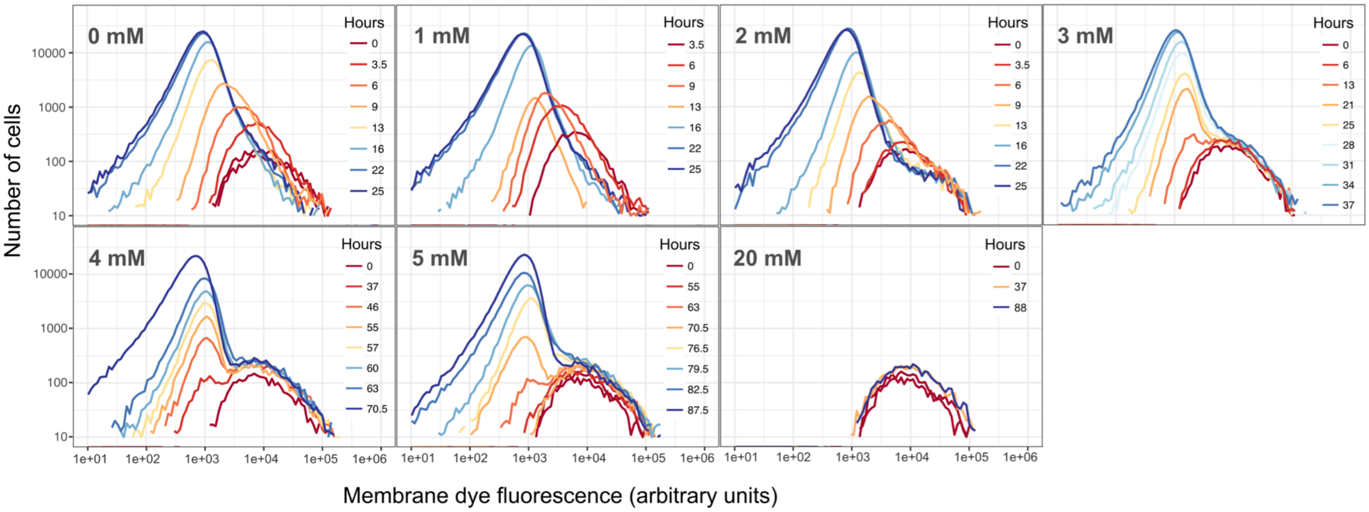


**Figure S5. Cell proliferation assays support the hypothesis that growth of *M. extorquens* in the presence of formaldehyde is due to a small subpopulation of tolerant cells, and that the abundance of tolerant cells decreases with increasing formaldehyde.**
(See also Fig. 3.) A naive culture of *M. extorquens* was stained using a fluorescent membrane dye and divided among culture flasks with different concentrations of formaldehyde; samples were periodically removed, fixed, and then analyzed by flow cytometry. Panels show formaldehyde treatment concentration; each line is the outline of a histogram of per-cell fluorescence measurements from one timepoint. For each sample, an equal volume was analyzed: higher cell counts indicate a higher density of cells in the culture. Per-cell membrane fluorescence decreases with growth, as cell division results in the dilution of membrane dye between daughter cells. Non-growing cells show no change in per-cell fluorescence. At 0 mM formaldehyde, all cells grow; at 20 mM, no cells grow; at intermediate concentrations, populations contain some growing and some non-growing cells.

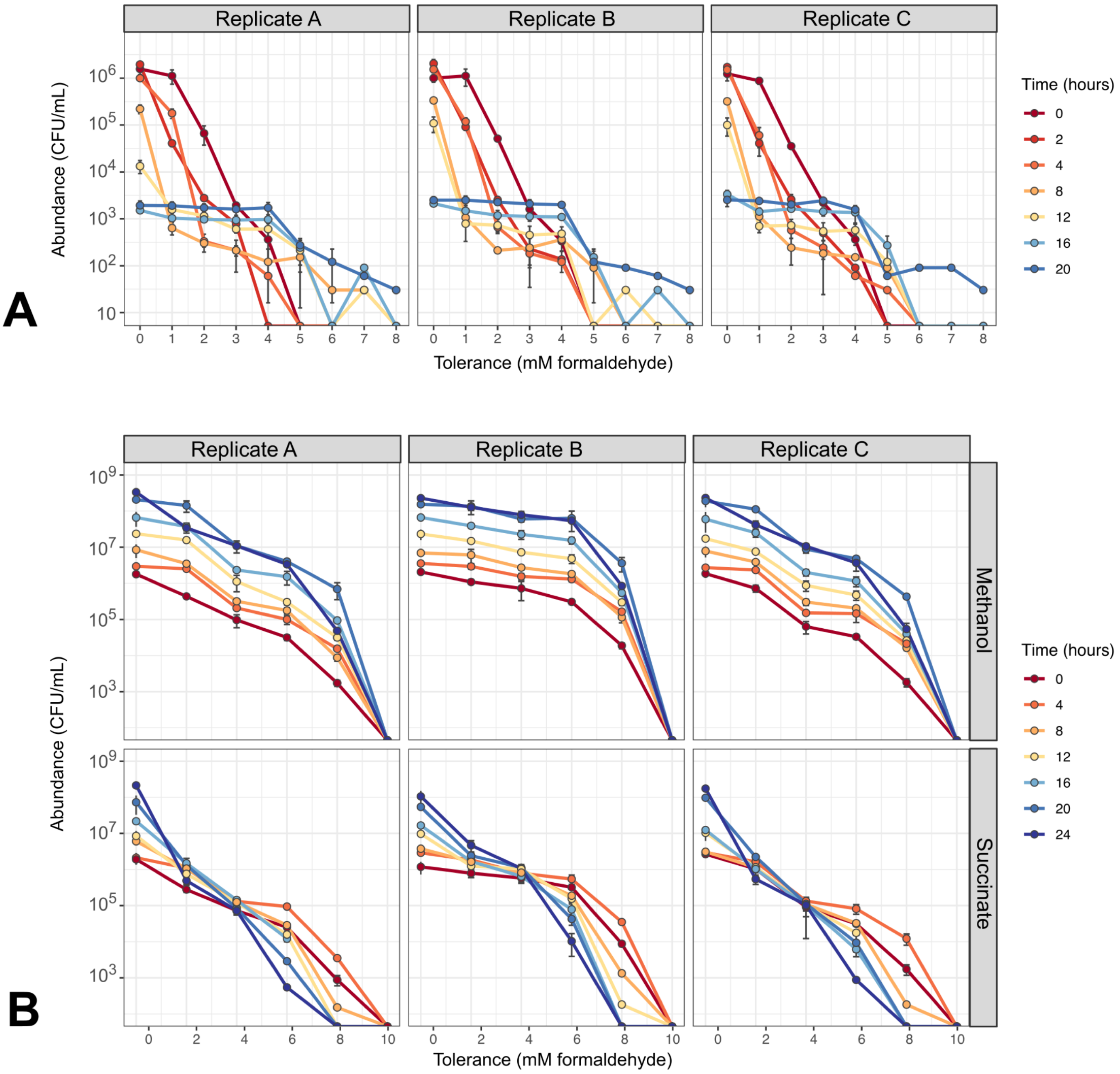


**Figure S6. The distribution of formaldehyde tolerance within an *M. extorquens* population changes over time depending on growth conditions.**
A) exposure to 4 mM formaldehyde selecting for tolerant cells; and B) regrowth of a tolerant population on formaldehyde-free medium (top row: methanol as the growth substrate; bottom row: succinate). Each column represents a separate biological replicate; error bars denote the standard deviation of three replicate platings for colony counts. Replicate B alone is shown in Fig. 7.

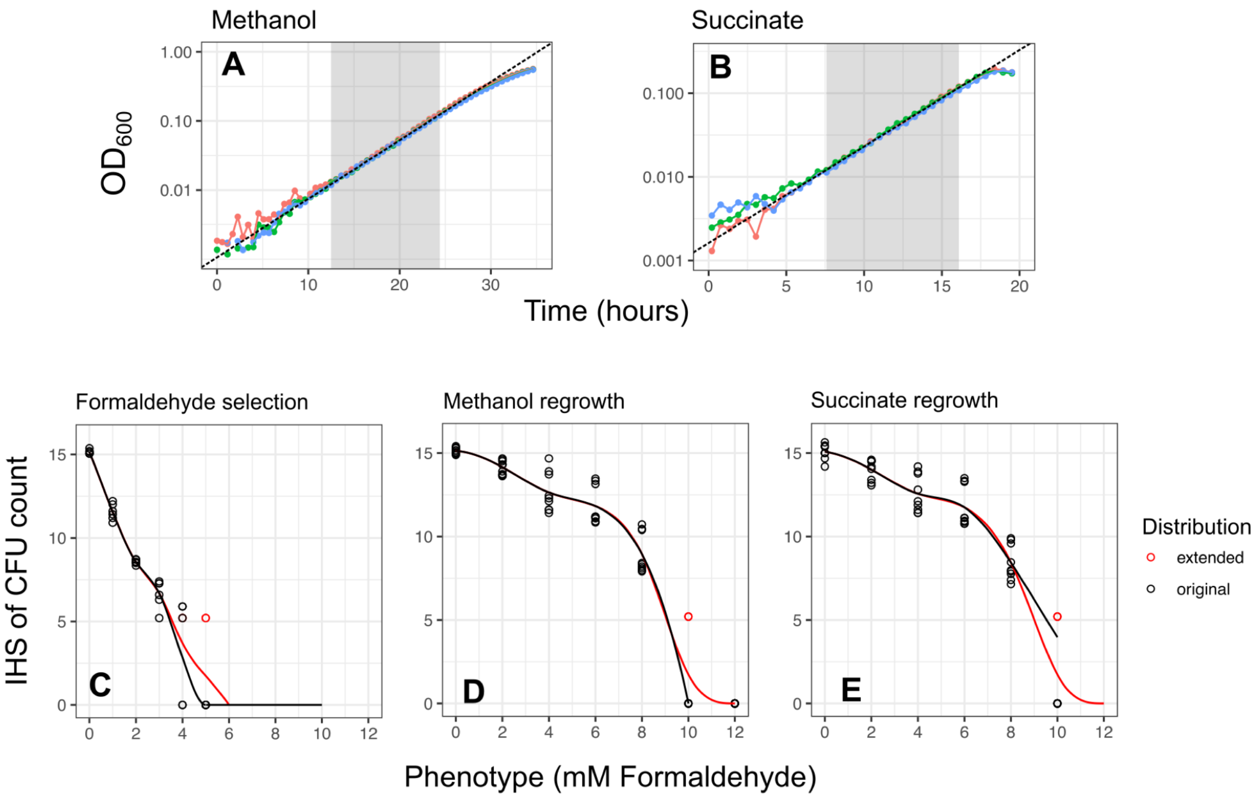


**Figure S7. Estimation of growth rates and initial conditions for use in the mathematical model.**
A) and B) Growth rates on methanol and succinate ($r_m$, $r_s$ respectively) were estimated by linear regression using data from *M. extorquens* cultures grown without formaldehyde on each of the two growth substrates. Gray shaded region indicates portion of growth curve used; dashed lines show exponential fit. $OD_{600}$: optical density at 600 nm. C-E) Initial conditions used for each of the three model scenarios. Points show experimental data (transformed by inverse hyperbolic sine) and line shows spline fit. Black points and line indicate original measured distribution; red points and lines indicate distribution extended to correct for undetected high-tolerance cells possibly present at abundances below the experimental detection limit. The extended distributions were used for the model.

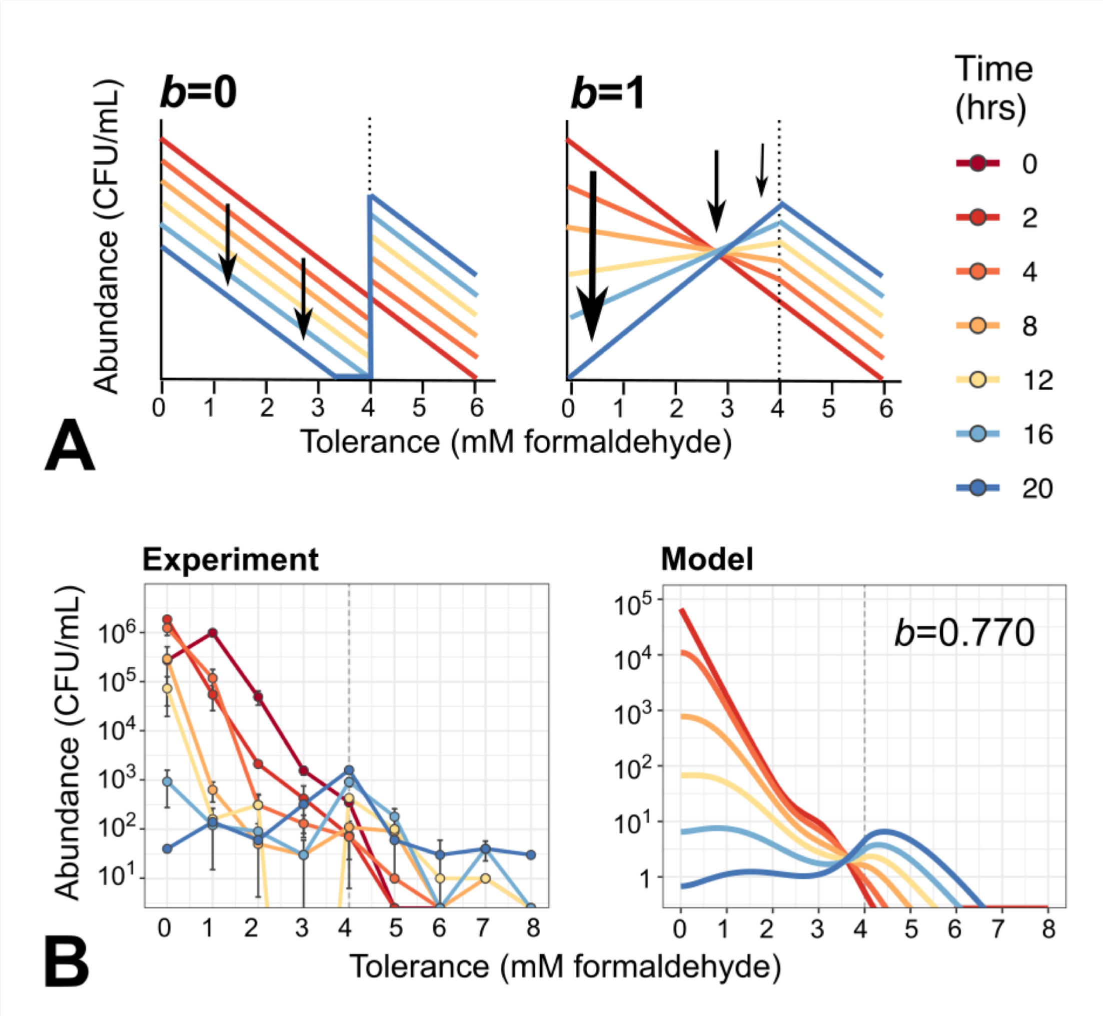


**Figure S8. The parameter *b* (dependence of death rate on formaldehyde tolerance) determines the shape of the population's phenotypic tolerance distribution after exposure to formaldehyde.**
Distributions are displayed in the non-cumulative form (see Methods for details). A) Schematic showing theoretical population shifts at exposure to 4 mM formaldehyde. When *b*=0, all cells die at the same rate regardless of their formaldehyde tolerance, as long as their tolerance level is lower than the formaldehyde concentration. In this case, formaldehyde exposure results in a tolerance distribution with two peaks: at $x$=0 and $x$=$F$. When *b*=1, death rate is proportional to tolerance, such that cells with higher $x$ die more slowly. This results in a tolerance distribution with one peak: at $x$=$F$. B) Results of (left) experiment and (right) model simulation: distribution falls partly between the two extremes. Note that the experimental results shown here are transformed into the non-cumulative form for comparison, as described in the Methods; and CFU abundances in model appear lower than in experiment because the continuous results have not been binned into 1-mM increments for this display.

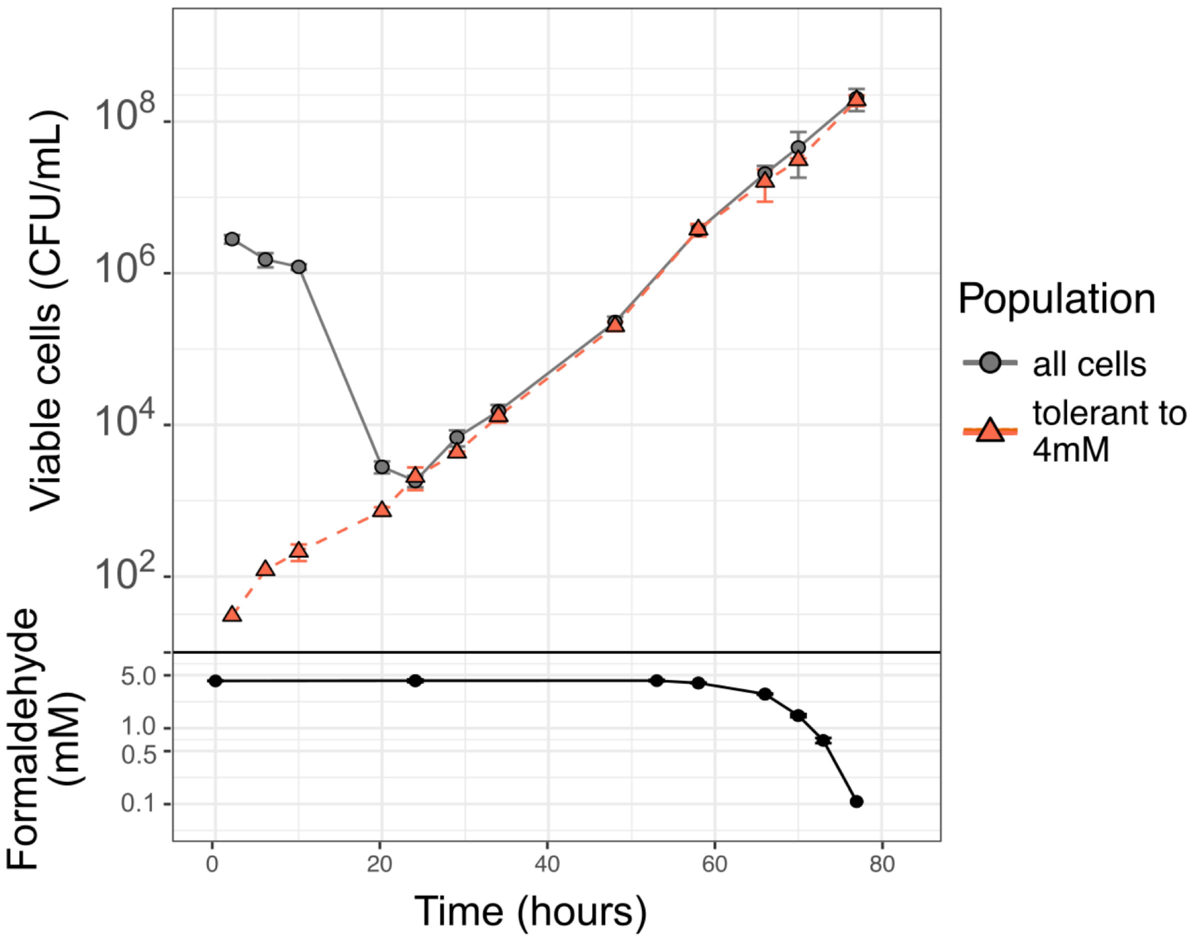


**Figure S9. Cells expressing mCherry show the same formaldehyde tolerance heterogeneity as wild-type *M. extorquens* cells.**
To facilitate identification of cells by flow cytometry in the cell proliferation assay, we used CM3839, a strain of *M. extorquens* PA1 engineered for the constitutive expression of the red fluorescent protein mCherry [64]. To ensure that the red-fluorescent strain showed the same phenotypic heterogeneity in formaldehyde tolerance as the wild-type strain (CM2730), we conducted a 4 mM formaldehyde exposure experiment with CM3839. Just as with CM2730 (Fig. 2), CM3839 populations contain a small subpopulation of cells that are tolerant to 4 mM formaldehyde, and that grow at a normal rate when the population is exposed to formaldehyde at that concentration. Error bars denote the standard deviation of three replicate measurements (for the formaldehyde measurements, error bars fall within the area of the symbols).

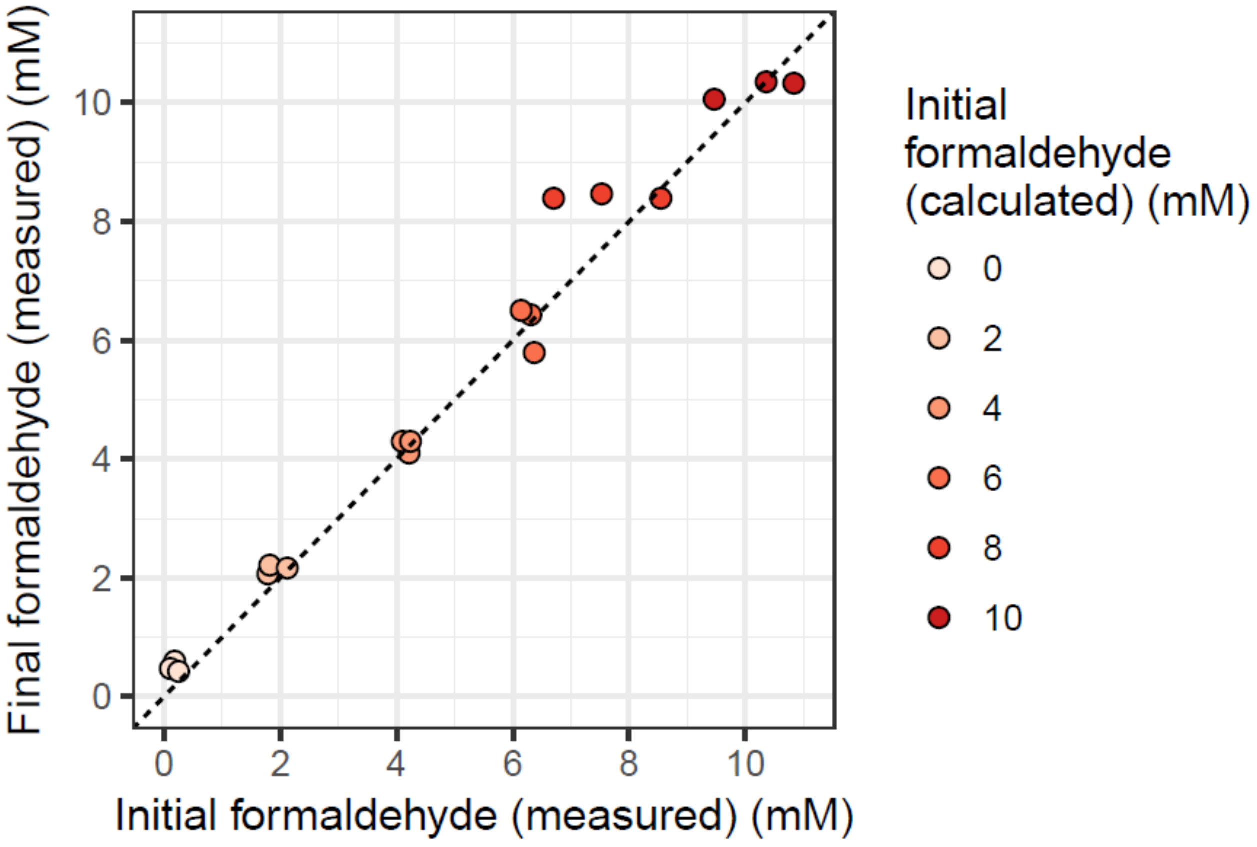


**Figure S10. Formaldehyde concentrations in agar growth medium are stable over time and reflective of similar concentrations in liquid medium.**
MPIPES-methanol-agar culture plates were made with the indicated concentration of formaldehyde. A small amount of agar (~0.1 g) was excised from the plate, melted, diluted 1:10 in MPIPES medium, and assayed for formaldehyde as described in Methods. Each plate was assayed in triplicate. Plates were then incubated for 3 days, stored together in the same bag, at 30ºC, and assayed again. No significant change in concentration was detected in any of the plates.

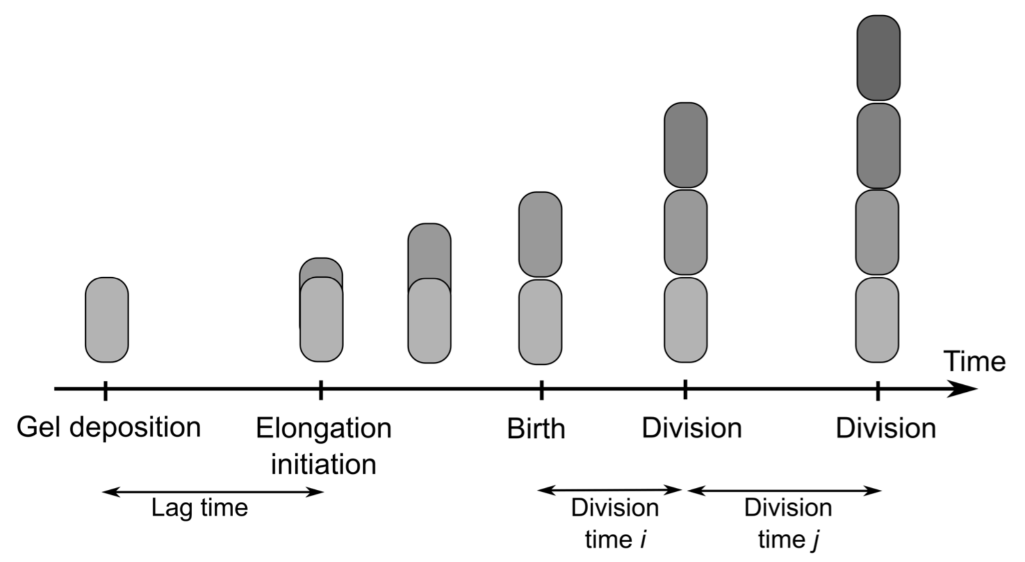


**Figure S11. Time-lapse microscopy: cell segmentation and tracking.**
Colony lag times and cell division times were measured as shown above: “lag time” refers to the time between deposition of cells in the gel and the beginning of cell elongation; there is only 1 lag time for each microcolony. “Division time” refers to the full time period between the formation of a cell and the formation of its daughter cell; for each microcolony, many division times were observed.

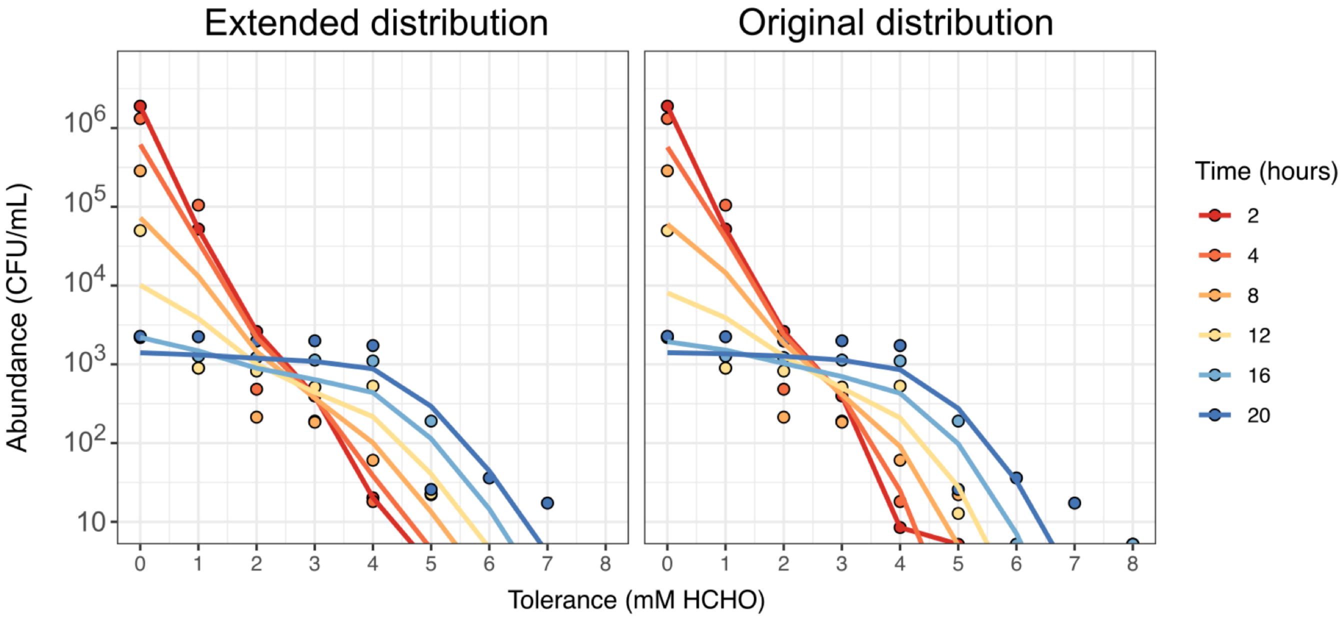


**Figure S12. Models using extended and original tolerance distributions perform similarly.**
Both panels show results of model simulation of population shifts during 4 mM formaldehyde selection experiment. Points: experimental data used for fitting model (averages of all replicates). Lines: model output, binned at 1 mM increments and summed to form cumulative distributions, for comparison with experimental data. Left panel: dataset in which extra CFU counts have been added at a few high-tolerance phenotype levels according to the rules described in the Methods, in order to correct for cells potentially present at abundances below the limit of detection (as in Fig. 9; included here for comparison). Right panel: dataset using only the experimentally measured counts, without extension.

**Table S1. Results of model selection using original data set for fitting (distribution not extended to account for experimental limit of detection).**
See Table 1 for comparison. For the likelihood ratio test, the name of the model used as the null, as well as the $\chi^2$ value and *p*-value, are given. Gray shading: the best-supported model. Pseudo-$R^2$ value for that model: 0.970.

| ***Model*** | ***Experimental scenario*** | | ***Parameters*** | | | | ***Likelihood Ratio Test*** | | |
|---|---|---|---|---|---|---|---|---|---|
| | **Condition** | **Substrate** | ***α*** | ***b*** | ***v*** | ***D*** | **null model** | $\chi^2$ | ***p*** |
| **F1** | selection | Methanol + Formaldehyde | 0.141 | n/a | n/a | n/a | n/a | n/a | n/a |
| **F2a** | selection | Methanol + Formaldehyde | 0.186 | 0.925 | n/a | n/a | F1 | 5.710 | 0.017 |
| **F2b** | selection | Methanol + Formaldehyde | 0.166 | n/a | -0.077 | n/a | F1 | 40.026 | <0.001 |
| **F2c** | selection | Methanol + Formaldehyde | 0.158 | n/a | n/a | 0.041 | F1 | 40.853 | <0.001 |
| **F3a** | selection | Methanol + Formaldehyde | 0.214 | 0.809 | n/a | 0.027 | F2b | 9.379 | 0.002 |
| **F3b** | selection | Methanol + Formaldehyde | 0.163 | n/a | -0.047 | 0.017 | F2b | 1.647 | 0.199 |
| **F4** | selection | Methanol + Formaldehyde | 0.220 | 0.929 | 0.051 | 0.044 | F3a | 0.752 | 0.386 |
